# Mechanism of an animal toxin-antidote system

**DOI:** 10.1101/2024.06.11.598564

**Authors:** Lews Caro, Aguan D. Wei, Christopher A. Thomas, Galen Posch, Ahmet Uremis, Michaela C. Franzi, Sarah J. Abell, Andrew H. Laszlo, Jens H. Gundlach, Jan-Marino Ramirez, Michael Ailion

## Abstract

Toxin-antidote systems are selfish genetic elements composed of a linked toxin and antidote. The *peel-1 zeel-1* toxin-antidote system in *C. elegans* consists of a transmembrane toxin protein PEEL-1 which acts cell autonomously to kill cells. Here we investigate the molecular mechanism of PEEL-1 toxicity. We find that PEEL-1 requires a small membrane protein, PMPL-1, for toxicity. Together, PEEL-1 and PMPL-1 are sufficient for toxicity in a heterologous system, HEK293T cells, and cause cell swelling and increased cell permeability to monovalent cations. Using purified proteins, we show that PEEL-1 and PMPL-1 allow ion flux through lipid bilayers and generate currents which resemble ion channel gating. Our work suggests that PEEL-1 kills cells by co-opting PMPL-1 and creating a cation channel.

## Main Text

Selfish genetic elements ensure their inheritance, even at the expense of host fitness. Toxin-antidote (TA) systems are one class of selfish element, made up of genetically linked toxin and antidote genes. The toxin is cytoplasmically inherited across generations while the cognate antidote is expressed in progeny. In animal TA systems, the parental toxin is transmitted to progeny via the sperm or egg. Offspring that do not inherit the TA genetic element are affected by the toxin, typically resulting in death or developmental defects (*1*), whereas the offspring which inherit the TA genetic element express the cognate antidote to prevent toxicity. Therefore, TAs guarantee their presence in the next generation by killing non-inheriting offspring. Recent work suggests that TA elements are more common among animals than previously thought (*2, 3*). However, the mechanisms of toxin and antidote activity remain largely unknown for animal TA systems.

One of the best characterized animal TA systems is *peel-1/zeel-1* in *C. elegans* (*4*). The *peel-1* toxin is expressed during sperm development but is not toxic to sperm. Mature sperm carry PEEL-1 protein and fertilization delivers the toxin to embryos, resulting in developmental arrest (*5*). However, offspring which inherit the TA element express the antidote *zeel-1* and do not arrest (Fig. 1A). Expression of PEEL-1 in adult worms also causes death (*5*), suggesting that toxicity is not specific to a particular developmental stage. Furthermore, PEEL-1 acts cell-autonomously in *C. elegans*; ectopic expression of PEEL-1 in specific tissues causes death of those cells, with no defects in neighboring cells (*5*). So far, no adult somatic cells have been found to be immune to PEEL-1, suggesting that toxicity works by disrupting a fundamental cellular process. In this study, we dissect the molecular mechanism of PEEL-1 toxicity. We find that PEEL-1 co-opts a conserved membrane protein of unknown function, PMPL-1. Together these proteins are sufficient to create a toxic, cation leak channel. Thus, our work determines the molecular mechanism of toxicity of an animal toxin-antidote system.

**Fig.1.**
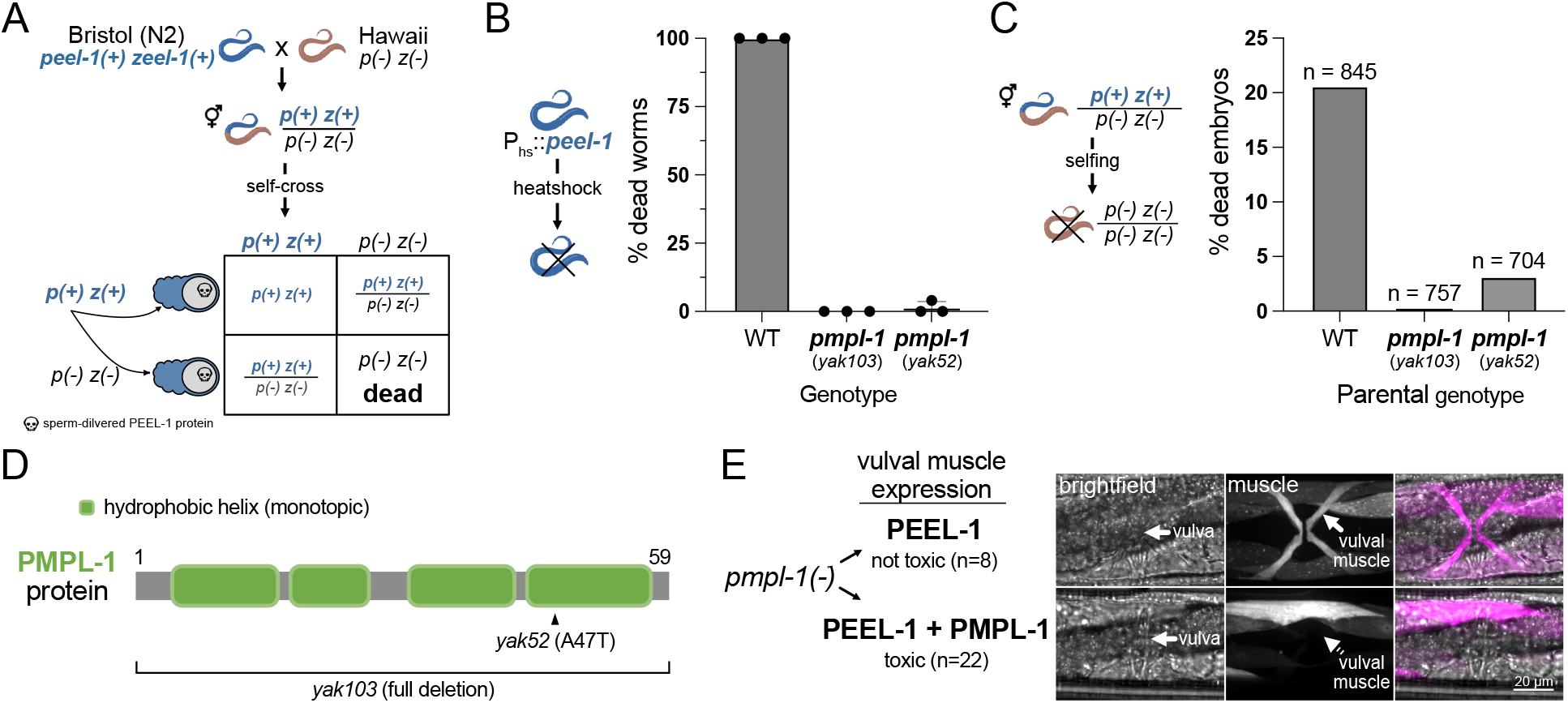
PMPL-1 is necessary for PEEL-1 toxicity in *C. elegans*. **(A)** Selfish activity of the *peel-1 zeel-1* toxin-antidote system in *C. elegans* shown in a genetic cross of strains from Bristol (which has the genetic element) and Hawaii (which lacks the genetic element). Hermaphrodite worms heterozygous for the presence of *peel-1 zeel-1 (p(*+*) z(*+*) / p(-) z(-))* have 25% inviable progeny. This is due to sperm-delivered PEEL-1 toxicity causing developmental arrest of *zeel-1(-)* progeny. **(B)** Proportion of worms dying from ectopic, heat shock-PEEL-1 expression. Two *pmpl-1* mutant alleles (*yak103* and *yak52*) provide resistance to toxicity. **(C)** Proportion of dead, arrested embryos from self-fertilizing hermaphrodites heterozygous for *peel-1 zeel-1*. n = total progeny scored. **(D)** Predicted domain structure of the PMPL-1 protein with mutant alleles shown. The hydrophobic helices are predicted to be monotopic, passing through one leaflet of a lipid bilayer. **(E)** Body wall and vulval muscle (magenta) of *pmpl-1*(*yak103*) worms with vulval muscle-specific expression of PEEL-1 alone (top) or PEEL-1 and PMPL-1 (bottom). Vulval muscles appears normal with PEEL-1 alone but are missing or atrophied when PEEL-1 and PMPL-1 are co-expressed in these cells. All worms expressing PEEL-1 and PMPL-1 in vulval muscle cells had missing or severely deformed vulval muscles (n=22). *peel-1* and *pmpl-1* are both GFP tagged. All channels are shown in fig. S2D. Scale bar = 20μm.

## Results

### PMPL-1 is required for PEEL-1 toxicity

To identify other genes required for PEEL-1 toxicity, we performed a large forward genetic screen for PEEL-1 suppressors in *C. elegans*. We mutagenized worms carrying transgenes for heat-shock inducible *peel-1* expression (*hsp-16*.*41p*::*peel-1*) (*5*). We isolated only two full-suppressors of heat-shock PEEL-1 toxicity, and both suppressors carry mutations in F47B7.1 (hereafter named *pmpl-1*) (Fig. 1B). Endogenous, sperm-delivered PEEL-1 is also suppressed by *pmpl-1* mutations (Fig. 1C), but not via a paternal-effect (fig. S1), suggesting that PMPL-1 acts in embryos to facilitate PEEL-1 toxicity and does not act in sperm. *pmpl-1* codes for a 59 amino acid protein predicted to be an integral membrane protein (Fig. 1D). The *pmpl-1* mutants were identified as a missense allele (*yak52*, A47T) and a full-gene deletion allele (*yak103*) (see Materials and Methods).

The *pmpl-1* expression pattern is consistent with its role in PEEL-1 toxicity. The *pmpl-1* promoter drives expression in embryos prior to toxicity from sperm-delivered PEEL-1 (fig. S2A) (*5*). *pmpl-1* is also widely expressed in several adult tissues (fig. S2B), consistent with heat-shock PEEL-1 toxicity in adults (*5*). Publicly available RNA-seq data (*6*) indicates that *pmpl-1* expression levels are lowest in the male gonad compared to all other tissues, consistent with wild-type sperm being unaffected by PEEL-1 (fig. S2C). These data suggest that PMPL-1 is required for cell susceptibility to PEEL-1 toxicity. We further confirmed that co-expression of PEEL-1 and PMPL-1 in the vulval muscle cells of *pmpl-1* mutants caused specific toxicity in this tissue (Fig. 1E and fig. S2D). Defects were not observed in surrounding tissues, indicating that PEEL-1 and PMPL-1 act in the same cell to cause toxicity.

### PMPL-1 is a conserved membrane protein of unknown function

PMPL-1 belongs to the Plasma Membrane Proteolipid 3 (PMP3) family of proteins, so we named it “PMP3-Like protein 1.” PMP3 proteins are widely present in bacteria, plants, and fungi, and are found in some simple animals (*7*). The role of PMP3 proteins in animals is unknown, but PMP3 proteins in plants, fungi, and bacteria are important for cold-stress resistance, membrane protein trafficking, and ion homeostasis (*8*–*11*). *pmpl-1* mutant worms do not have any obvious phenotypes. However, PMPL-1 is highly conserved among nematodes (fig. S3). We identified 15 PMP3-like proteins in *C. elegans* (fig. S4) through BLAST searches. PMPL-2 (also known as TXT-9) is most similar to PMPL-1 (75% similarity) and was found in an RNAi screen for defective transcellular chaperone signaling (*12*). However, there is no known molecular function for any *C. elegans* PMP3-like protein.

PMP3 proteins are thought to contain two transmembrane spanning domains (*13, 14*), consistent with DeepTMHMM predictions for PMPL-1 (fig. S5A) (*15*). However, AlphaFold2 predicts PMPL-1 as a monotopic protein, with four helices passing through only one leaflet of a lipid bilayer (Fig. 1D and fig. S5B) (*16, 17*). We favor the monotopic prediction of PMPL-1 because two recently solved structures of a bacterial photosynthetic complex showed a PMP3 protein within the complex having a similar monotopic structure (*18, 19*).

In contrast to *pmpl-1*’s broad conservation in nematodes (fig. S3), *peel-1* is found only in *C. elegans* and has no homology to any known protein (*5*). Additionally, *pmpl-1* is genetically unlinked from *peel-1 zeel-1*. These features suggests that PMPL-1 has biological roles other than supporting PEEL-1 toxicity and that PEEL-1 co-opts PMPL-1 for its own use.

### PEEL-1 and PMPL-1 are sufficient for toxicity in HEK293T cells

Given that *pmpl-1* was the only full suppressor of PEEL-1 toxicity found in our screen, we hypothesized that PEEL-1 and PMPL-1 may be the only two components required for toxicity. We expressed these proteins in human embryonic kidney cells (HEK293T) and assayed for cytotoxicity using lactate dehydrogenase (LDH) release into the culture media as a measure of plasma membrane rupture (*20*). We found that each protein alone was not toxic, but co-expression of PEEL-1 and PMPL-1 resulted in significant cytotoxicity (Fig. 2A). Other PMP3-like proteins such as *C. elegans* PMPL-2 and yeast PMP3 were not toxic with PEEL-1 (Fig. 2B), suggesting that PMPL-1 has a specific role in toxicity that is not universal to the PMP3 family. The reconstitution of toxin activity in a heterologous system suggests that PEEL-1 and PMPL-1 are sufficient for toxicity and may act by disrupting an essential, conserved cellular process.Antidote activity could also be reconstituted in HEK293T cells by co-expression of ZEEL-1, resulting in reduced toxicity (Fig. 2A).

**Fig.2.**
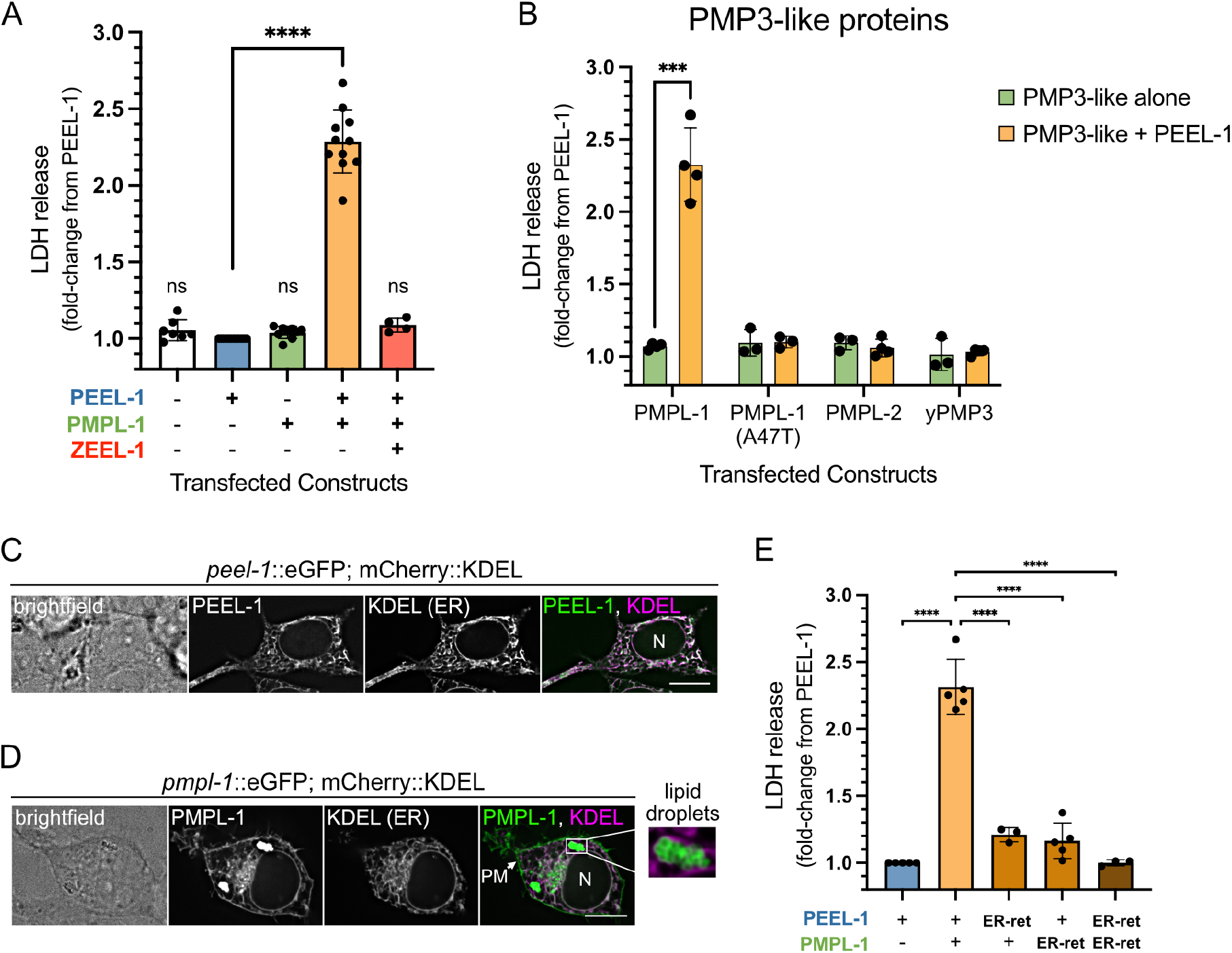
PEEL-1 and PMPL-1 are sufficient for toxicity in HEK293T cells. **(A)** Cytotoxicity (measured by LDH release) of combinations of three constructs transfected in HEK293T cells. Each data point is a biological replicate, normalized to LDH release from transfection with *peel-1*::eGFP in the same experiment. All plots show means with SD. Transfections combine constructs encoding for mCherry or eGFP (-) or a fluorescent-tagged protein (+): PEEL-1::eGFP (top), PMPL-1::mCherry (middle), or mCherry::ZEEL-1 (bottom). **(B)** Cytotoxicity of PMP3-like proteins alone or with PEEL-1::eGFP. The PMPL-1 *yak52* (A47T) mutant protein, *C. elegans* PMPL-2, and the yeast homolog yPMP3 are shown. **(C)** Live-cell imaging of a single cell transfected with an ER-marker (mCherry::KDEL) and *peel-1*::eGFP or **(D)** *pmpl-1*::eGFP. The cell nucleus is indicated (N). PMPL-1 is also seen on the plasma membrane (PM) and on lipid droplets (inset). Scale bar = 10μm. **(E)** Cytotoxicity is suppressed by addition of the GBR1, ER-retention tag on the C-terminus of PEEL-1::eGFP or PMPL-1::mCherry. P-values in (A) and (E) calculated using one-way ANOVA with Dunnett’s multiple comparisons test, comparing all samples to PEEL-1 alone in (A) and all samples to PEEL-1 with PMPL-1 in (E). In (B), multiple unpaired t-tests were used with Holm-Šídák test, comparing each PMP3-like protein alone to PMP3-like with PEEL-1 (***, *p* < 0.001; ****, *p* < 0.0001).

### Plasma membrane localization of PEEL-1 and PMPL-1 is critical for toxicity

PEEL-1 and PMPL-1 localize to several membrane-bound compartments in HEK293T cells and *C. elegans*. Both proteins localize to the endoplasmic reticulum (ER), while PMPL-1 also localizes to the plasma membrane (PM) and to lipid droplets (Fig. 2C-D and fig. S6A), consistent with the predicted monotopic topology. Although PEEL-1::eGFP is not easily detectable on the PM of transfected HEK293T cells, stable expression of PEEL-1::eGFP in HEK293 cells or PEEL-1::tagRFP in *pmpl-1* knock-out worms shows clear PM localization (fig. S6B-C), suggesting that PEEL-1 does localize to the plasma membrane and does not require PMPL-1 for this localization.

To determine in which compartment these proteins perform their toxic roles, we prevented their movement to the PM by the addition of a GBR1 ER-retention tag (*21*). ER-retention tags on either protein resulted in more than an 80% drop in toxicity, and retention tags on both proteins completely suppressed toxicity (Fig. 2E), suggesting that PEEL-1 and PMPL-1 do not act in the ER and likely perform their toxic roles at the PM.

### PEEL-1 toxicity causes cell swelling in HEK293T cells

LDH release is an end-point measure of plasma membrane rupture, but it was unclear whether plasma membrane rupture was a primary or secondary effect of toxicity. Imaging 48 hours after transfection showed that 92% of HEK293T cells co-expressing PEEL-1 and PMPL-1 were swollen, about double their typical size (Fig. 3A-B). Live imaging showed cells exhibiting abnormal phenotypes approximately 16 hours after transfection. Cells began with slight swelling of the nucleus and cell, followed by jetting out round protrusions of 5-10 μm, similar in size to the nucleus (Fig. 3C and movie S1). These large protrusions are short-lived, typically existing for less than 10 minutes before being reabsorbed by the cell. Eventually, cells swell evenly in all directions (Fig. 3D). Swollen cells often lose the integrity of their plasma membrane, followed by ER fragmentation and ER swelling (fig. S7). By 48 hours after transfection, we observe various outcomes for swollen cells: cell lysis, cell detachment from the plate, and swollen, intact cells attached to the plate. Our LDH assay likely captures only the first of these phenotypes and may underestimate the fraction of cells experiencing toxicity. The cell swelling phenotype is reminiscent of the necrotic vacuoles previously described in PEEL-1 affected embryos in *C. elegans* (*5*).

**Fig.3.**
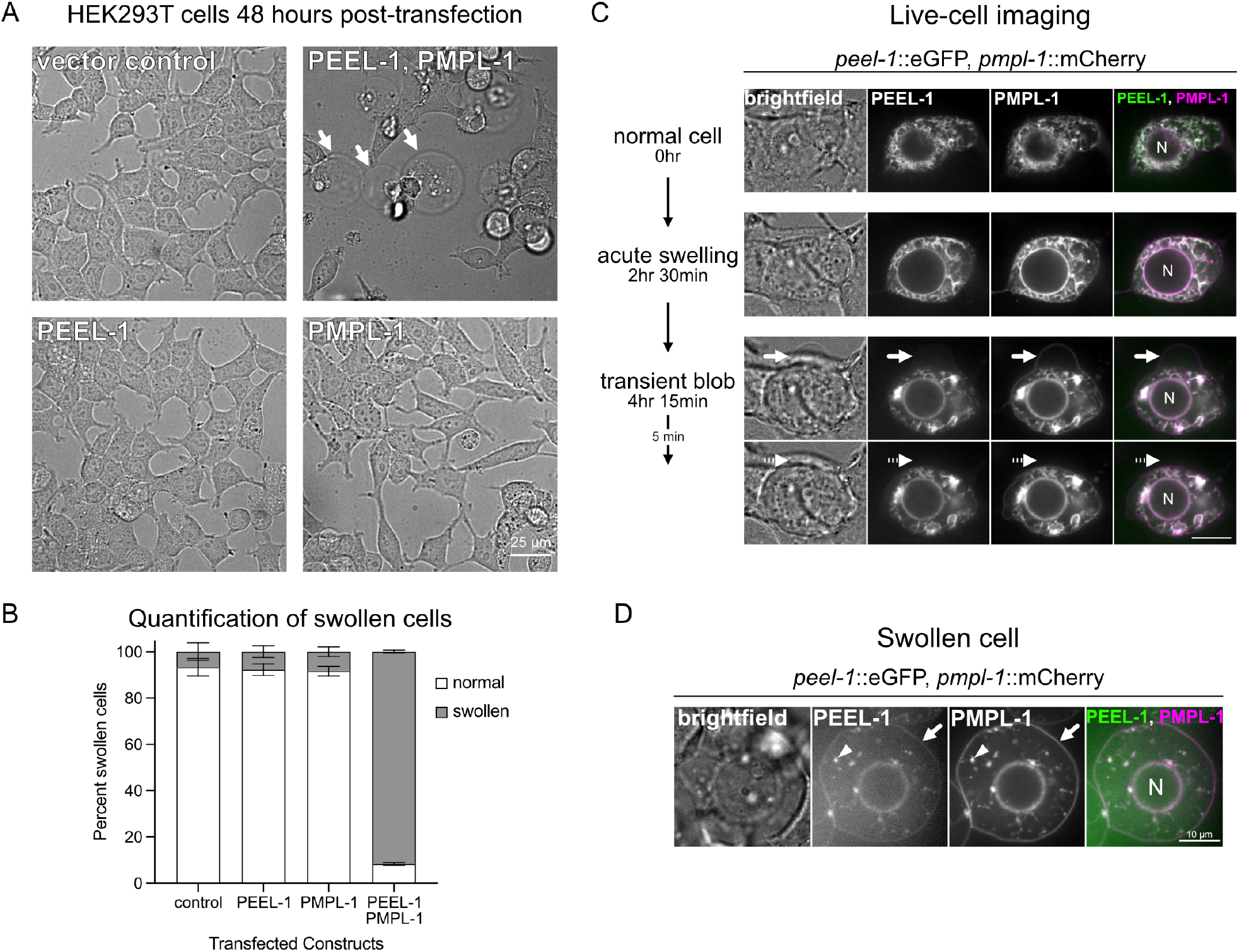
PEEL-1 toxicity results in HEK293T cell swelling. **(A)** Brightfield images of cells transfected with indicated constructs. Cells were imaged 48 hours after transfection. Scale bar = 25 μm. Arrows point to swollen cells. **(B)** Percent of transfected cells that appear normal or swollen under brightfield. Mean with SD of three biological replicates is shown. 100 cells were scored for each condition in each biological replicate. **(C)** Selected frames of a live-cell imaging time-course experiment (see movie S1). A single cell is shown, co-expressing PEEL-1::eGFP and PMPL-1::mCherry. The nucleus is labeled (“N”). t=0 is 16 hours after transfection. Acute swelling can be seen (t=2 hr 30min), followed by a transient blob or protrusion (arrow) jetted out by the cell (t=4 hr 15min) and later reabsorbed (dotted arrow) (t=4 hr 20min). Scale bar = 10 μm. **(D)** A typical phenotype from a cell transfected with PEEL-1::eGFP and PMPL-1::mCherry at 48 hours post-transfection. PEEL-1 and PMPL-1 can be seen on the plasma membrane (arrow) and in fragmented ER (arrowhead). Scale bar = 10 μm.

The cell swelling phenotype caused by PEEL-1 toxicity suggests that cell death is non-apoptotic (*22*). Indeed, *C. elegans* mutants defective in apoptosis (*ced-3*) and apoptotic cell engulfment (*ced-2* and *ced-5*) are still susceptible to PEEL-1 toxicity (fig. S8) (*23*–*25*). Other mammalian cell death pathways such as pyroptosis and necroptosis appear to be absent from *C. elegans* (*26*). Thus, the cytotoxic phenotypes we observe are likely due to a direct effect of PEEL-1 and PMPL-1 activity rather than indirect induction of programmed cell death.

### PEEL-1 has an amphipathic helix that is critical for toxicity

PEEL-1 has no homology to known proteins (*5*), so we turned to structural predictions to identify important regions in PEEL-1. AlphaFold2 predicted a low-confidence structure with six alpha-helices (Fig. 4A) (*16, 17*). The longest four helices matched DeepTMHMM’s prediction of four transmembrane domains (fig. S5C). Closer inspection revealed that the fourth predicted transmembrane domain is amphipathic (Fig. 4B). An amphipathic helix (AH) has opposing hydrophilic and hydrophobic faces and is a critical feature of known pore-forming toxins. For example, actinoporins are a diverse family of toxins which oligomerize their AHs to construct channels (*27*). The properties of the predicted PEEL-1 AH are similar to actinoporin AHs. The PEEL-1 AH is 22 amino acids in length (residues 111-132), making it long enough to span the lipid bilayer (*28*). The hydrophobic moment (μH), a measure of the degree of amphipathicity, is within the range of actinoporin channel-forming toxins (fig. S9) (*29*). One notable difference is that the PEEL-1 AH is more hydrophobic than actinoporin AHs (fig. S9), suggesting that the PEEL-1 AH may always exist within the membrane, unlike in actinoporins which have soluble AH conformations.

**Fig.4.**
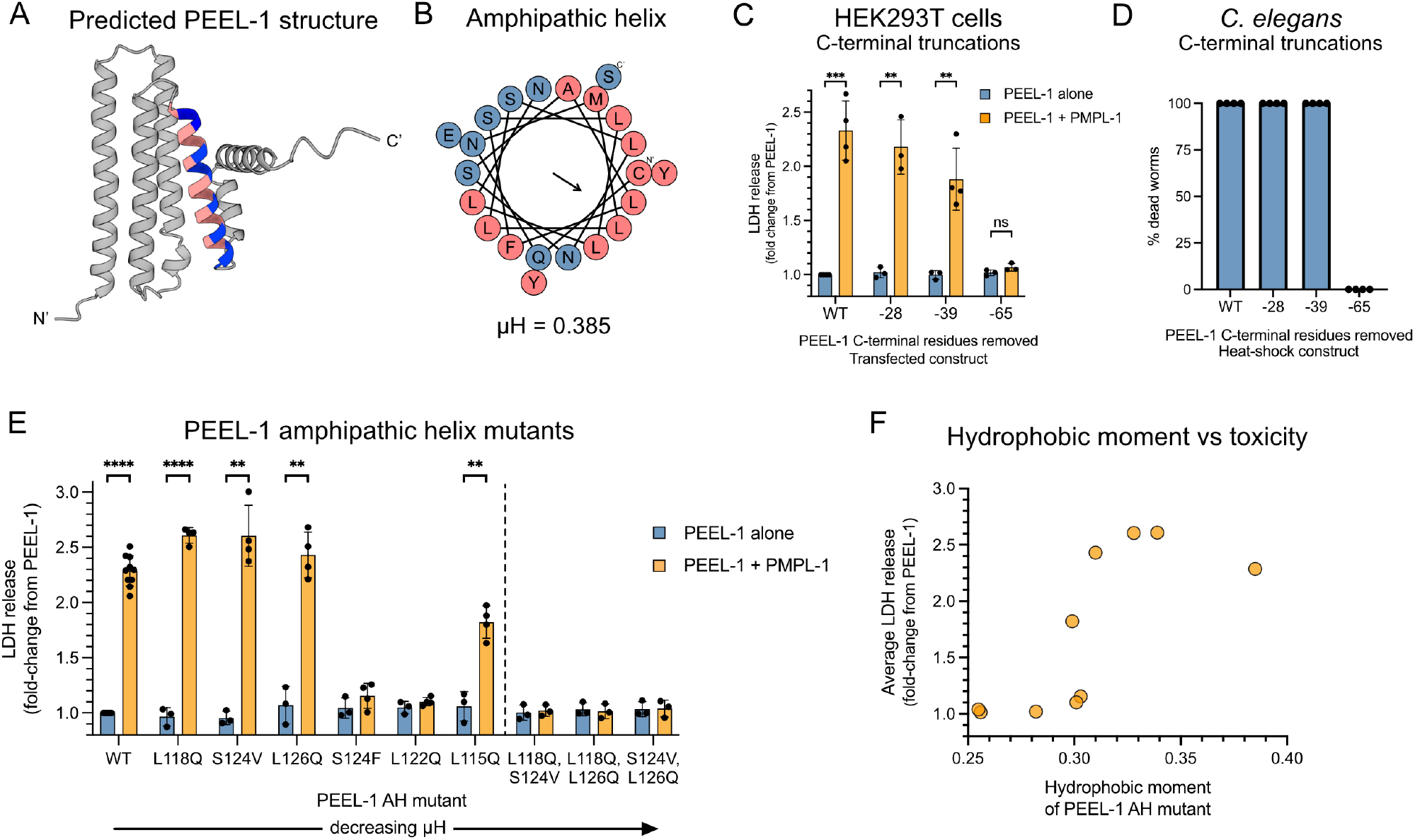
PEEL-1’s amphipathic helix is critical for toxicity. **(A)** The AlphaFold2 predicted structure of PEEL-1 and **(B)** a helical wheel representation of the putative PEEL-1 amphipathic helix. Amphipathic helix residues are colored (pink = hydrophobic, blue = hydrophilic). **(C)** Cytotoxicity of a series of PEEL-1 C-terminal truncations expressed in HEK293T cells. The number of amino acids removed are indicated (ex. “-28” means the last 28 residues were removed). Each truncation removes an additional alpha helix. The “-65” truncation removes the amphipathic helix. **(D)** Percent dead worms after heat-shock PEEL-1 expression of the indicated truncation mutant. 50 worms were assayed for each data point, and two independent transgenic lines were tested for each construct. **(E)** Cytotoxicity of PEEL-1 amphipathic helix missense mutants in HEK293T cells. Mutants are ordered by descending hydrophobic moment (μH). Six single mutants (left of dotted line) and three double mutants are shown (right of dotted line). All bar graphs show mean with SD. Statistics were performed using multiple unpaired t-tests with Holm-Šídák test, comparing each PEEL-1 alone to PEEL-1 and PMPL-1 (**, *p* < 0.01; ***, *p* < 0.001; ****, *p* < 0.0001). **(F)** The average toxicity of missense mutants from (E) plotted against their hydrophobic moment.

We hypothesized that the PEEL-1 AH may perform a similar role as it does in actinoporins, where AHs construct the lining of a toxic channel. To test this, we determined whether this helix was required for toxicity in HEK293T cells and *C. elegans*. We found that deleting the last two helices of PEEL-1 did not severely impair toxicity (“-28 aa” and “-39aa” in Fig. 4C-D). However, deleting the AH resulted in complete loss of toxicity in both mammalian cells and worms (“-65 aa” in Fig. 4C-D). By deleting one amino acid at a time, we determined that toxicity in HEK293T cells was eliminated after removing at least 40 amino acids (fig. S10). This PEEL-1(-40aa) mutant contains only three residues following the AH, possibly destabilizing the AH.

We also tested whether the amphipathic property of the PEEL-1 AH was critical for toxicity by creating a series of missense mutants that modified the AH’s hydrophobic moment. Six single-missense mutants were tested. Three of these mutants attenuated PEEL-1 toxicity (S124F, L122Q, L115Q) while three mutants were still fully toxic (L118Q, S124V, L126Q) (Fig. 4E). However, all pairwise combinations of these three fully toxic mutants resulted in complete loss of toxicity (Fig. 4E). Across the nine missense mutants we tested, toxicity correlated well with the amphipathicity of the AH (Fig. 4F). These data suggest that the amphipathic property of the PEEL-1 amphipathic helix is critical for toxicity, supporting its potential role as the lining of a channel. However, we cannot rule-out alternative explanations for the importance of the AH, including roles in protein-protein interaction or protein stability.

### PEEL-1 and PMPL-1 create a non-specific monovalent cation channel

We hypothesized that PEEL-1 toxicity was caused by disruption of ionic gradients across the cell membrane, leading to osmotic imbalance and subsequent cell swelling. This could be through two mechanisms: an ion leak channel, where specific ions flow down their electrochemical gradient, or a non-selective pore, causing flux of all ions and smaller osmolytes across the plasma membrane. To test these possibilities, we performed whole-cell patch-clamp electrophysiology on HEK293 cells in physiological ion conditions (high internal K^+^, high external Na^+^). Currents from cells transfected with *peel-1* or *pmpl-1* alone appeared similar to currents from untransfected cells (Fig. 5A and fig. S11A-B). Obtaining recordings of cells co-transfected with *peel-1* and *pmpl-1* was difficult, due to variable timing of toxicity, and was complicated by loss of cells due to swelling. Therefore, we generated tetracycline-inducible cell lines that more tightly regulate PEEL-1 and PMPL-1 expression. We generated a HEK293 cell line with tetracycline-inducible PMPL-1::mCherry and then introduced stable expression of either eGFP (“Control” cell line) or PEEL-1::eGFP (“Experimental” cell line). Cells with and without tetracycline are hereafter referred to as induced and uninduced, respectively. Nearly all Experimental cells swell 24 hours after induction (fig. S12A), with the first signs of swelling occurring 6 hours after induction (fig. S12B).

**Fig.5.**
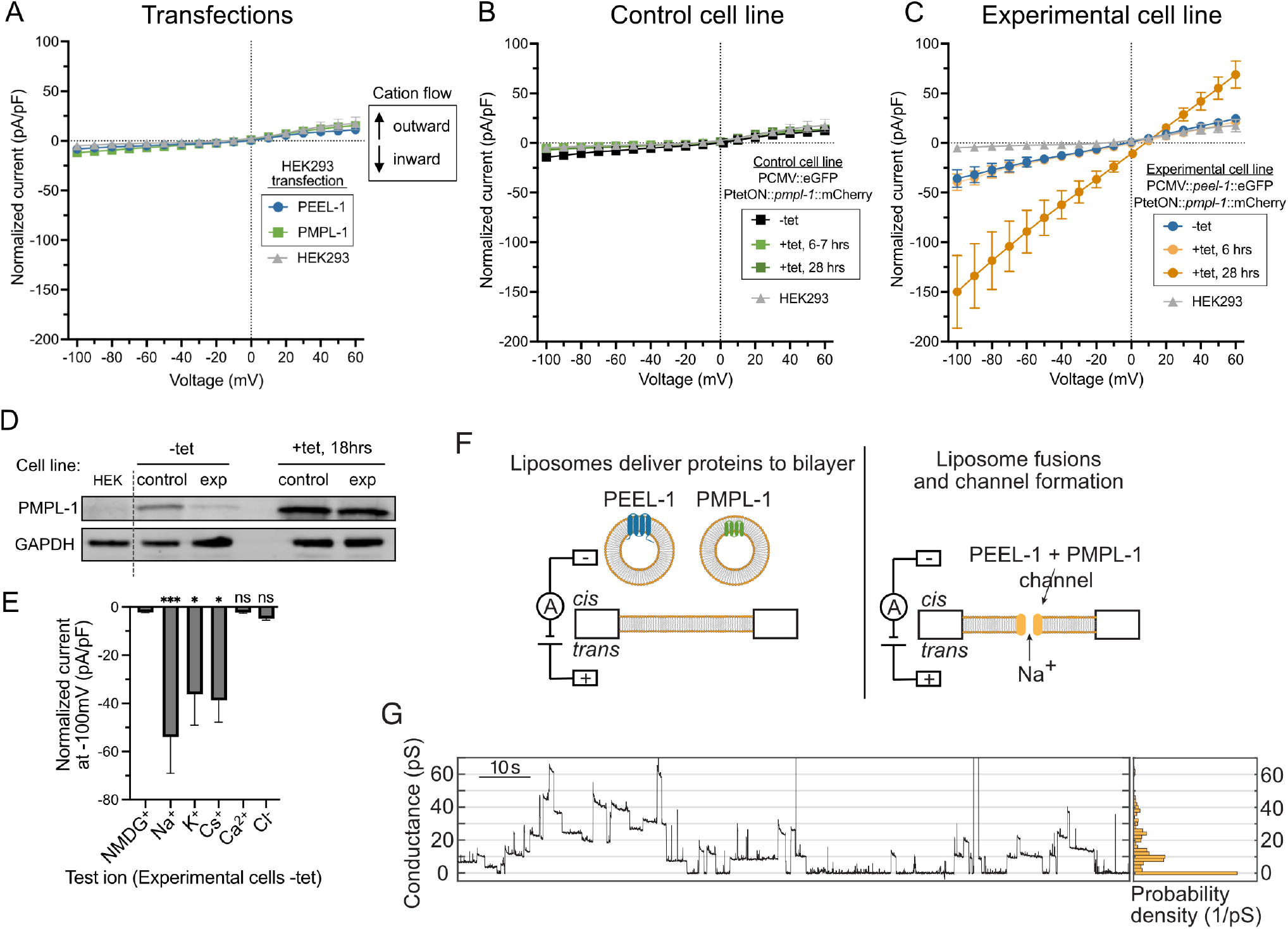
PEEL-1 and PMPL-1 create a monovalent cation channel. Current-voltage plots of whole-cell patch-clamp electrophysiology on **(A)** transfected cells, **(B)** Control cell line, and **(C)** Experimental cell line. High intracellular potassium (140 mM K^+^/ 8.6 mM Na^+^) and high extracellular sodium (145 mM Na^+^/ 4 mM K^+^) solutions are used. Currents elicited by a family of 0.5 second voltage steps from a -30 mV holding potential, from -100 mV to 60 mV, in 10 mV increments. Currents normalized to cell capacitance (pF). Negative and positive currents indicate cation flow into or out of the cell, respectively. **(A)** Plots from HEK293 cells acutely transfected with *peel-1*::eGFP or *pmpl-1*::mCherry. **(B)** Plots from Control cell line expressing constitutive eGFP and tetracycline-inducible *pmpl-1*::mCherry. **(C)** Plots from Experimental cell line expressing constitutive *peel-1*::eGFP and tetracycline-inducible *pmpl-1*::mCherry. Control and Experimental cell lines are shown without tetracycline or with tetracycline at the indicated time after addition of tetracycline. All plots shown as mean with SEM. **(D)** Western blot for PMPL-1::mCherry in Control and Experimental cell lines without tetracycline (-tet, left) and 18 hours after addition of tetracycline (+tet, right). Leaky expression of PMPL-1 is seen in both cell lines in the absence of tetracycline. Less background PMPL-1 expression is seen in Experimental cells than in Control cells, likely because of selection against higher background PMPL-1 expression when in combination with PEEL-1 but not eGFP. GAPDH loading control shown. **(E)** Permeability of indicated ions was assayed in Experimental cells without tetracycline. Test ionic solutions substituted previous bath solution (145 mM Na^+^/ 4 mM K^+^) with 140 mM pure cations (external) or 20 mM anions (internal), except Ca^2+^ (20 mM external, 1 mM internal Ca^2+^ with 2.5 mM EGTA). All plots show mean with SEM. Statistical tests compare all results to NMDG (treated as control) in one-way ANOVA with Dunnett’s multiple comparisons test (*, *p* < 0.05; ***, *p* < 0.001). **(F)** Schematic of planar lipid bilayer experiment. Liposomes containing purified PEEL-1 or PMPL-1 are added to the *cis* well to deliver proteins to the lipid bilayer. **(G)** Conductance traces of one experiment (left) and a histogram of the trace (right, 2 pS bin width, normalized based on probability density) at an applied voltage of +180 mV. SDS-PAGE gels of purified proteins are shown in fig. S15A and more example conductance traces and controls are shown in fig. S16.

We performed whole-cell patch clamp recordings without leak subtraction of uninduced cells, and at 6-7 hours and 28 hours after induction (Fig. 5B-C). Similar to untransfected naïve HEK293 cells, uninduced Control cells exhibited small instantaneous negative (inward) currents at negative potentials, and these currents did not increase after induction (Fig. 5B and fig. S11C). In contrast, uninduced Experimental cells exhibited a larger instantaneous inward current compared to uninduced Control cells and naïve HEK293 cells, and the magnitude of these currents further increased after 28 hours of induction (Fig. 5C and fig. S11C). When clamped at the most negative potential (-100 mV), inward currents from uninduced and 28-hour induced Experimental cells were 7-fold and 30-fold greater than naïve HEK293 cells, respectively. These data suggest that co-expression of PEEL-1 and PMPL-1 creates a constitutively active ionic channel.

The inward currents observed in uninduced Experimental cells were instantaneous, exhibiting no voltage-dependent gating kinetics (Fig. 5C and fig. S11C), consistent with a leak channel constitutively open at physiological resting membrane potentials. This ion leakage in uninduced Experimental cells was surprising since these cells appear healthy and do not have obvious growth defects. We detected some PMPL-1 expression in uninduced Control and Experimental cells (Fig. 5D and fig. S12A), consistent with previous reports of leaky background expression from tetracycline-inducible promoters (*30*–*32*). Therefore, we reasoned that uninduced Experimental cells may have sufficient background PMPL-1 expression to display an electrophysiological phenotype but not enough to cause cell swelling or cell death.

To characterize the ionic selectivity of channels formed by PEEL-1 and PMPL-1 co-expression, we performed ion substitution experiments. We took advantage of the inward currents generated by uninduced Experimental cells since these cells are morphologically normal, reducing the possibility of contamination by endogenous HEK293 channels secondarily activated by cell swelling or cell death such as the LRRC8-associated volume-regulated anion channels (VRACs) (*33, 34*). We substituted the physiological mixed cation bath solution (high Na^+^/low K^+^) with equivalent bath solutions containing single cations (Na^+^, K^+^, Cs^+^, NMDG^+^ or a combination of 20 mM Ca^2+^/120 mM NMDG^+^) and monitored instantaneous inward currents at negative potentials. We found that bath solutions with Na^+^, K^+^, or Cs^+^ still yielded large inward currents, while bath solutions with NMDG^+^ and Ca^2+^ abolished all inward currents (Fig. 5E and fig. S13). These results suggest that PEEL-1 and PMPL-1 co-expression creates a channel that is permeable to monovalent cations (Na^+^, K^+^, and Cs^+^) but impermeant to the divalent Ca^2+^ cation and the bulkier cation NMDG^+^. We next tested for Cl^-^ permeability by recording with high internal Cl^-^ (140 mM KCl) and low external Cl^-^ (140 mM NMDG^+^). Any Cl^-^ permeability at negative potentials would cause Cl^-^ movement out of the cell and be recorded as a negative (inward) current. We observed no inward currents (Fig. 5E and fig. S13), indicating that the channel is impermeable to Cl^-^. Altogether, these results suggest that PEEL-1 and PMPL-1 create an ion channel rather than a non-selective pore. The PEEL-1 and PMPL-1 channel conducts monovalent cations and is impermeable to anions and divalent cations. An increase in PMPL-1 expression in induced Experimental cells may result in excessive influx of Na^+^, sufficient to overwhelm compensatory volume regulatory mechanisms, leading to osmotic dysregulation, cell swelling, and cell death.

### Predicted PEEL-1 pentamer structure has features of an ion channel

Ion channels are typically made up of oligomeric complexes, so we used AlphaFold2 to determine possible oligomeric structures of PEEL-1. Structural predictions were of low-confidence due to PEEL-1’s lack of homology to known proteins (*5*). Nevertheless, the predicted PEEL-1 pentamer has a striking resemblance to known cation channels (fig. S14A-D and data S1): (a) the outside surface of the complex is hydrophobic, consistent with its location in lipid bilayers, (b) the complex has a central, uninterrupted hydrophilic pore, providing a potential path for ions through the complex, and (c) the opening of the pore region is surrounded by a ring of negatively charged residues. These features are strikingly similar to the structure of the ZAR1 cation channel, which is a toxic pentameric channel with a ring of acidic residues at the extracellular mouth of the pore that serve as a cation selectivity filter (*35*). In the predicted PEEL-1 pentamer, a similar ring of negative charges is formed by five D109 residues. Mutating this residue to an alanine (D109A) resulted in complete loss of PEEL-1 toxicity (fig. S14E). This residue is two amino acids before the PEEL-1 AH. Five copies of the PEEL-1 AH meet in the middle of the complex, forming the pore-like region of the predicted structure (fig. S14A-B), consistent with our model of PEEL-1 AHs constructing the lining of a channel.

### *In vitro* reconstitution of the PEEL-1/PMPL-1 ion channel

We used an independent approach to test if PEEL-1 and PMPL-1 create an ion channel by assaying whether these purified proteins allow ions to flow through artificial planar lipid bilayers (Fig. 5F). PEEL-1 and PMPL-1 were individually expressed in *E. coli* with maltose-binding protein (MBP) tags, affinity purified in detergent, and incorporated into liposomes for delivery to planar lipid bilayers (fig. S15A and Materials and Methods). For PEEL-1, the full-length protein was purified and incorporated into liposomes, along with co-purified truncation products (fig. S15A). After adding PEEL-1 or PMPL-1 liposomes to the planar lipid bilayer, we observed no currents (fig. S16B-C). After adding both PEEL-1 and PMPL-1 liposomes to the same bilayer, we observed discrete, step-like conductance events similar to those created by typical ion channels (Fig. 5G and fig. S16D-E) (*35*–*38*). There was variation in single channel conductance and open duration (Fig. 5G and fig. S16D-E). Such variability could reflect heterogenous stoichiometries, or it could be due to technical limitations such as misfolded proteins, truncated PEEL-1 proteins in the sample, mixed orientation of proteins in the bilayer, or imprecise amounts of each protein in the bilayer. Nevertheless, the stepwise conductance *in vitro* and cation-selective property in cells suggest that PEEL-1 and PMPL-1 together construct a monovalent cation channel which causes osmotic dysregulation and cell death.

## Discussion

Our study provides unprecedented understanding of an animal TA system. We provide evidence that the PEEL-1 toxin co-opts a conserved membrane protein to create a cation channel which ultimately causes cell swelling and death. We show that PEEL-1 may be the primary structural component of the channel and hypothesize that PMPL-1 is required to gate the channel open. PEEL-1 may have evolved this co-option mechanism for temporal control over ion channel activity, to avoid toxicity in sperm and early embryos. Prevention of off-target toxicity in the germline and before the maternal-to-zygotic transcriptional switch is likely an evolutionary hurdle faced by many animal TA systems. Therefore, co-option may be a common evolutionary solution to prevent off-target toxicity in animal TA systems.

## Materials and methods

### Worm strains and maintenance

Worm strains were maintained using standard procedures (*39*). Strains used in this study are provided in table S1.

### Isolation of suppressors of heat-shock induced PEEL-1

Worm strains XZ1047 and XZ1372 were mutagenized using ENU or EMS (*39, 40*). These strains contain two copies of *hsp-16*.*41p*::*peel-1* in order to avoid isolating suppressors that have mutations in the transgene. F2 populations of worms were heat-shocked at 34°C for 2 hours and allowed to recover at room temperature overnight. Surviving worms were isolated. Approximately 200,000 haploid genomes were screened. *yak52* was isolated from EMS mutagenesis of XZ1047, while *yak103* was isolated from ENU mutagenesis of XZ1372.

We isolated a total of six independent suppressors of heat-shock PEEL-1 toxicity. Four of the mutations were partial suppressors while two were full suppressors. The causative genes were identified using genetic mapping and whole-genome sequencing. The four partial suppressors have mutations in mRNA export factors *nxf-1* and *nxt-1*, but act by affecting expression of *hsp-16*.*41p::peel-1* and do not suppress toxicity of endogenous, sperm-delivered PEEL-1 (*41*).

The two full suppressors *yak52* and *yak103* are recessive suppressors of PEEL-1 toxicity and have no other obvious phenotype. They were determined to be allelic by complementation testing and were mapped to the X chromosome by crossing to strains carrying visible or fluorescent markers on each chromosome (strains EG1000, EG1020, EG8040, EG8041). The closest linkage seen was to the visible marker *lon-2*.

### Identification of *pmpl-1*

Strains XZ1177 and XZ1307 carrying 4X and 5X outcrossed *pmpl-1(yak52)* and *pmpl-1(yak103)* were subjected to whole-genome sequencing. DNA was purified according to the Hobert laboratory protocol (http://hobertlab.org/whole-genome-sequencing/). Illumina (XZ1177) or Ion Torrent sequencing (XZ1307) was performed at the University of Utah DNA Sequencing Core Facility. The dataset for XZ1177 contained approximately 17,000,000 reads of a mean read length of 36 bases, resulting in ∼6X average coverage of the *C. elegans* genome. The dataset for XZ1307 contained approximately 12,800,000 reads of a mean read length of 147 bp, resulting in ∼19X average coverage. The sequencing data were processed on the Galaxy server at usegalaxy.org (*42*). SNPs and indels were identified and annotated using the Unified Genotyper and SnpEff tools (*43, 44*). Although we found genes on the X chromosome in each strain containing nonsynonymous mutations, none of these genes were affected independently in both strains. Thus, to identify potential deletions, we used BEDtools Genome Coverage tool to calculate sequencing coverage across the genome in intervals of consecutive bases with the same coverage (*45*). Intervals with zero sequencing coverage were then annotated using SnpEff, filtered to intervals on the X chromosome which overlapped protein-coding regions, and sorted by length. In XZ1307, the largest of these intervals on the X chromosome was found near where *yak103* and *yak52* had been mapped and was subsequently confirmed to correspond to *yak103*, a 323-bp deletion spanning the F47B7.1 gene, from its 5’UTR to 3’UTR and deleting all the coding sequence. In XZ1177, we found two small regions of F47B7.1 lacking coverage in our whole genome sequencing. Via Sanger sequencing, we identified *yak52* as a nonsynonymous mutation in one of these regions, with a G to A mutation that results in an A47T substitution.

### Heat-shock PEEL-1 toxicity assay

Heat-shock PEEL-1 assays were performed with 50 gravid adults of each strain for each biological replicate. Adults were picked to NGM plates seeded with OP50 and heat-shocked (2 hours at 34°C). After recovering at room temperature for 2 hours, moving worms were counted. Non-moving worms were tested for response to touch stimulus and scored as dead if they did not move.

### Sperm-delivered PEEL-1 toxicity assay

Sperm-delivered PEEL-1 toxicity in wild-type or *pmpl-1* mutant backgrounds was assayed by counting unhatched embryos. To make *pmpl-1*; *peel-1(*+*) zeel-1(*+*) / peel-1(-) zeel-1(-)* worms (Fig. 1C), AFS216 *peel-1(-) zeel-1(-)* males were crossed to *peel-1(*+*) zeel-1(*+) worms with *pmpl-1* mutations *yak103* (XZ2194), *yak52* (XZ2283), or wild-type *pmpl-1* (N2, control). F1 males were backcrossed to the parent strain (XZ2194, XZ2283, or N2) to obtain the desired genotype. To determine if *pmpl-1* acts by a paternal effect (fig. S1), F1 males were backcrossed instead to AFS216 *peel-1(-) zeel-1(-)*.

Embryonic lethality was assayed on individual, self-fertilizing hermaphrodites. Single worms were allowed to lay embryos for 16-24 hours before being removed from the plate. Plates were left for one more day to allow embryos to develop and hatch, and the progeny were scored. Unhatched embryos and larval worms were counted. Unhatched embryos were scored as dead. Progeny were genotyped in bulk for *zeel-1(-)* by PCR using oGP65 (5’attctggagttcgtgaggtgc3’) and oGP66 (5’ccctcctttcccacccaac3’). Only plates showing *zeel-1(-)* alleles were considered to have parents with the desired genotype, heterozygous *peel-1(*+*) zeel-1(*+*) / peel-1(-) zeel-1(-)*.

### Plasmid construction

All plasmids used in this study are provided in table S2. Plasmids used for mammalian cell transfection were generated using Gibson assembly (*46*) using N1 vector backbone (CMV promoter). Constructs for tetracycline-inducible expression were cloned into pFTSH vector (gift from Nancy Maizels). Plasmids used for *C. elegans* transgenics were constructed using either Gibson assembly or Gateway cloning (Invitrogen). Typical Gateway cloning was performed using a 3-fragment approach (promoter, coding region, and fluorophore-UTR or UTR) into a destination vector, pCFJ150. Restriction digest was used to confirm the plasmid was correct. Sanger sequencing was used to confirm that constructs did not contain mutations (table S2). Transgenic worms were made using microinjection. For each transgenic strain, we isolated at least two independent lines and confirmed similar results.

### Microscopy

All microscopy was performed on a NikonTi2-E Crest X-light V2 spinning-disk confocal microscope. For live-cell imaging, mammalian cells were maintained in a humidified, heated chamber (37°C) in 5% carbon dioxide. For long-term imaging, the objective heater was also heated to 37°C. For imaging *C. elegans*, worms or embryos were picked onto a 2% agar pad, immobilized in a humidified chamber for 10 minutes in 33 mM sodium azide, and then immediately imaged.

### Mammalian cell maintenance and generation of clonal cell lines

All mammalian cells were grown at 37°C and 5% carbon dioxide in DMEM with GlutaMAX (Thermo Fisher Scientific), 10% fetal bovine serum (RMBIO or Gibco), and 100 U/mL of penicillin-streptomycin (Gibco). All transfection-based cytotoxicity experiments were done in HEK293T cells (gift from Mary Claire-King). For electrophysiology experiments, HEK293 cells (CRL-1573; ATCC) were used. Tetracycline-inducible cells (tetON cells) were made in HEK293 Flp-In TReX cells (Invitrogen) (gift from Nancy Maizels). tetON cells were grown in media made with Tet system approved fetal bovine serum (Gibco).

Stable tetON lines were generated by co-transfection of FlpO (pOG44; Life Technologies) and PtetON::*pmpl-1*::mCherry (pLC79, in pFTSH backbone, a gift from Nancy Maizels). Two days after transfection, cells were plated at low density in 10-cm plates with hygromycin B (150 μg/mL). Two clonal populations were picked into media with hygromycin B (150 μg/mL), grown to 80% confluency, and frozen in 10% DMSO at -80°C. One clonal population was used to generate the Experimental and Control cell lines.

Experimental tetON and Control tetON cell lines were generated in parallel. One tetON::*pmpl-1*::mCherry clonal cell line was transfected with the respective plasmid, *peel-* 1::eGFP (pGP9) or eGFP. Two days after transfection, cells were plated at low density in selective media containing both hygromycin B (150 μg/mL) and G418 (400 μg/mL). After about two weeks, multiple single colonies were picked and continued to be grown under selection. For the Experimental cell line, more than half the picked clones which grew in selective media did not have visible green fluorescence, presumably due to leaky *pmpl-1*::mCherry imposing selection against *peel-1*::eGFP-expressing cells. Therefore, only clones which had green fluorescence were maintained. This issue was not encountered during the generation of the Control cell line. All Experimental and Control cell lines were screened following tetracycline treatment. 48 hours after tetracycline treatment, all cell lines had red fluorescence, all Experimental cell lines had very few adhered cells (n= ∼6), and all Control cell lines had no obvious cellular phenotypes (n= ∼10).

### Cytotoxicity assay

Cytotoxicity in mammalian cells was measured using a colorimetric assay for LDH release (Promega CytoTox 96). 12-well plates were seeded with approximately 7×10^4^ HEK293T cells about 24 hours prior to transfection. Each well was transfected with a mixture of 1.5 μg DNA and 3 μg polyethylenimine (PEI) in 200 μL OptiMEM (Gibco). Three constructs were transfected in all experiments. For each of PEEL-1, PMPL-1, and ZEEL-1, we either transfected a fluorophore-tagged protein construct or the fluorophore-only vector as a control. Supernatant was collected 43-45 hours after transfection and used in 96-well plates for LDH assays, following manufacturer protocol. Two technical replicates were measured for each well and absorption at 490 nm was averaged between replicates. All experiments included wells for a non-killing control (fluorophore-encoding vectors), *peel-1*::eGFP alone, and *peel-1*::eGFP with *pmpl-1*::mCherry. The fold-change over *peel-1*::eGFP was calculated from transfections performed on the same day.

### SDS-PAGE and western blots

For lysis of mammalian cells, cells were collected from 12-well plates and washed with PBS. Cell pellets were lysed in radioimmunoprecipitation assay (RIPA) lysis buffer (25 mM Tris pH 7.4, 150 mM NaCl, 0.1% SDS, 1% NP-40, 1% sodium deoxycholate) with DNase and Halt Protease Inhibitor Cocktail (Thermo Fisher Scientific). After lysis (15 min on ice), the sample was centrifuged (21,000 x *g*, 10 min, 4°C), and the supernatant was used for SDS-PAGE.

Following separation by SDS-PAGE, proteins were transferred onto a nitrocellulose membrane. After blocking in Intercept Blocking Buffer (LI-COR, 1hr room temperature) membranes were incubated with primary antibody and nutated overnight at 4°C. Mouse monoclonal anti-mCherry (a gift from Jihong Bai and the Fred Hutch Cancer Center antibody development shared resource center; 1:1000), mouse monoclonal anti-MBP (New England Biolabs, E8032; 1:10,000), rabbit polyclonal anti-GAPDH (Sigma Aldrich, G9545; 1:5,000). Appropriate secondary antibodies were used, either goat anti-mouse or donkey anti-rabbit conjugated to Alexa 680 or Alexa 790 (Invitrogen). Membranes were imaged on an Odyssey CLx (LI-COR Biosciences).

### Electrophysiology

For electrophysiology on acutely transfected cells, plasmid constructs were transfected into HEK293 cells using Viafect reagent (Promega), following the manufacturer’s protocol. Cells were first prepared for transfection by plating onto 12-well tissue culture plates (Nunc 12-565-321; Thermo Fisher Scientific) at a density of ∼0.5-2 x10^5^ cells per well and grown to ∼80-90% confluence with standard media, allowing for one confluent well per transfection condition. On the day of transfection, media were replaced with 0.5 mL fresh DMEM with 10% fetal bovine serum (Gibco), without penicillin-streptomycin. Lipophilic/DNA transfection complexes were generated for each well, combining a total of ∼1.0 μg of plasmid DNAs with serum-free OptiMEM (Gibco) to a final volume of 100 μL, then adding 3.0 μL Viafect with gentle trituration, allowing the mixture to assemble at 24°C for 30 minutes, and then added to each well, dropwise. Transfected cells were incubated overnight at 37°C and visually monitored for transfection efficiency *in situ* using an inverted plate microscope equipped with fluorescence (Invitrogen EVOS M7000; Thermo Fisher Scientific). Transfection efficiencies were typically >70-80%. Specific amounts of plasmid DNAs for acute transfections per well: a) pCMV::*peel-1*::eGFP (0.8 μg), b) pCMV::*pmpl-1*::mCherry (0.1μg) and pcDNA3 (0.8 μg), c) pCMV::*peel-1*::eGFP (0.8 μg) and pCMV::*pmpl-1*::mCherry (0.1μg). Untransfected HEK293 cells served as controls.

Following overnight incubation, cells in transfected wells were dissociated with TrypLE (Gibco); and replated at low density onto 12 mm poly-*D*-lysine-coated glass coverslips (NeuVitro) in 24-well tissue culture plates (FisherBrand FB012929; Thermo Fisher Scientific), for patch-clamp electrophysiology. Typically, ∼10,000-15,000 cells were replated per well at sufficiently low density to isolate individual cells. This was necessary to prevent the formation of electrical junctions between contacting cells, which precludes adequate space-clamp recording conditions. Recordings were performed from 0.5 to 3 days after replating at low densities. Control and Experimental stable HEK293 cell lines were similarly replated at low density on coverslips for patch-clamp recordings.

For patch-clamp recordings, coverslips containing adherent cells were transferred to a Zeiss AxoExaminer.A1 microscope, equipped with a 40X water immersion objective and epifluorescence capability. Pipettes were positioned with a Sutter MPC-325 micromanipulator (Novato). Whole-cell voltage-clamp recordings were acquired with an AxoClamp200B amplifier (Molecular Devices), using pClamp10.4. Composition of recording solutions are listed below:

Bath Solutions:

1. *HEK293 bath (4 K*^+^, *145 Na*^+^*) (in mM)*: 4.0 KCl, 145 NaCl, 2.0 CaCl_2_, 2.0 MgSO_4_, 10 HEPES, 10 glucose, pH to 7.4 with NaOH.
2. *140 Na*^+^ *(in mM):* 140 NaCl, 2.0 CaCl_2_, 2.0 MgCl_2_, 10 HEPES, pH to 7.4 with NaOH.
3. *140 K*^+^ *(in mM):* 140 KCl, 2.0 CaCl_2_, 2.0 MgCl_2_, 10 HEPES, pH to 7.4 with KOH.
4. *140 Cs*^+^ *(in mM):* 140 CsCl, 2.0 CaCl_2_, 2.0 MgCl_2_, 10 HEPES, pH to 7.4 with CsOH.
5. *140 NMDG*^+^ *(in mM):* 140 N-methyl-*D*-glucamine (NMDG^+^), 2.0 CaCl_2_, 2.0 MgCl_2_, 10 HEPES, pH to 7.4 with HCl.
6. *20 Ca*^*2*+^ *(in mM):* 20 CaCl_2_, 120 NMDG^+^, 2.0 MgCl_2_, 10 HEPES, pH to 7.4 with HCl.

Internal Pipette Solutions:

1. *140 K*^+^, *low Cl*^*-*^ *(in mM)*: 140 K-*D*-gluconate, 1.0 CaCl_2_, 2.0 MgCl_2_, 10 HEPES, 2.4 EGTA, 4.0 Na_2_ATP, 0.3 Na_2_GTP, pH to 7.4 with KOH.
2. *140 K*^+^, *high Cl*^*-*^ *(in mM)*: 140 KCl, 1.0 CaCl_2_, 2.0 MgCl_2_, 10 HEPES, 2.4 EGTA, 4.0 Na_2_ATP, 0.3 Na_2_GTP, pH to 7.4 with KOH.

For all recordings of cationic currents, different bath solutions were used in combination with *140 K*^+^, *low Cl*^*-*^ internal pipette solution. For recordings of Cl^-^ currents, *140 NMDG*^+^ bath solution was used in combination with *140 K*^+^, *high Cl*^*-*^ internal pipette solution; under these conditions outward Cl^-^ conductance at negative potentials would be seen as inward currents by standard electrophysiological recording convention. Patch pipettes were pulled from borosilicate glass (1B120F-4; World Precision Instruments) on a P-97 Sutter Instruments puller (Novato), with resistances of 3.0-5.0 MΩ. Currents were allowed 3-5 mins to stabilize after achieving whole-cell recording configuration, filtered at 5 kHz, and acquired at 10 kHz. Series resistance compensation was >80% for all recordings.

Mean currents normalized to cell capacitance were analyzed and plotted using pClamp10.4 (Molecular Devices), Microsoft Excel, and OriginPro8.5 (Northampton). Leak subtraction correction was not applied to any of the data. Current-voltage (I/V) data were plotted as means with standard errors (SEMs) and statistical calculations were performed in Prism (GraphPad).

### Protein purification

MBP::PEEL-1::His_8_ and MBP::PMPL-1 were purified from *E. coli* BL21(DE3) and C43(DE3) cells, respectively. An overnight culture (37°C, 220rpm, 100 μg/mL ampicillin, 33 μg/mL chloramphenicol) was used to inoculate 1 L of LB media (100 μg/mL ampicillin, 220 rpm, baffled flask), grown at 37°C to OD_600_ of 0.4-0.7, and induced with IPTG (0.5 mM). PMPL-1 cells were chilled on ice prior to induction, and then induced in an 18°C shaker overnight. PEEL-1 cells were induced at 37°C for 1 hour since longer induction at lower temperatures resulted in decreased yields. Cells were pelleted and either stored at -80°C or used immediately for purification.

For purification, cell pellets were resuspended in Buffer A (HEPES pH 7.4, 150 mM NaCl, 5 mM 2-mercaptoethanol, 10% glycerol). Cells were lysed using a Dounce homogenizer, followed by 30 minutes rocking at room temperature after addition of DNase, lysozyme, protease cocktail inhibitor (Pierce), and n-octyl-β-glucopyranoside detergent (β-OG, 2% final concentration) (Anatrace). Lysate was clarified by centrifugation (20 min, 20,000 x *g*, 4°C) before batch binding with amylose resin for 1-2 hours at 4°C. Resin was pelleted (5 min, 700 x *g*) and resuspended in wash buffer to load onto a clean column. Four washes (5 column volumes) and four elutions (two column volumes) were performed by gravity flow. Wash buffer and elution buffer contained 1% β-OG in Buffer A. Maltose (10 mM) was included in elution buffer. Protein yield was estimated by absorbance at 260 nm. Proteins were stored at 4°C for less than three days before incorporation into liposomes.

### Proteo-liposomes

Liposomes containing MBP::PEEL-1::His_8_ or MBP::PMPL-1 were made with 50% DOPC:POPS (1,2-dioleoyl-sn-slycero-3-phosphocholine: 1-palmitoyl-2-oleoyl-sn-glycero-3-phospho-L-serine). Lipid stocks were purchased in chloroform (Avanti Polar Lipids) and β-OG stock was made in methanol. Lipid films were made by mixing lipids with β-OG detergent (lipid:detergent molar ratio of 4:35) in glass vials, dried under a nitrogen stream (10-20 min), and further dried in a Speedvac evaporator (4-6 hours). Buffer A was added to the dried lipid-detergent mixture and resuspended via two or three rounds of bath sonication (5 min, room-temperature) with nutation (20 min, 4°C). Protein was added in an approximate protein:lipid molar ratio of 1:400 (PMPL-1) or 1:2,500 (PEEL-1). Detergent was removed by dialysis: 500μL of sample was dialyzed (20 kDa cutoff) in 250 mL Buffer A with 0.5 g Biobeads SM-2 (Bio-Rad) for 16-18 hours at 4°C. Dialyzed sample was floated through a density gradient (Histodenz 35%, 25%, 0%; Sigma-Aldrich) using ultracentrifugation (SW60Ti, 55 krpm, 4°C, 30 min). Liposomes were collected from the top fraction (200 μL, using a wide-bore pipette tip), aliquoted, flash frozen in liquid nitrogen, and kept at -80°C. Liposomes were thawed fresh on the day of each experiment. A second liposome prep was made from an independent protein purification with two adjustments: soybean lipids (Avanti Polar Lipids) with cholesterol were used for liposomes instead of DOPC and POPS, and 1 mM EDTA was included in Buffer A.

### Synthetic planar lipid bilayers

Two ∼50 μL wells (*cis* and *trans*) linked by a ∼20 μm PTFE aperture were filled with Buffer B (20 mM HEPES, pH 7.4, 150 mM NaCl, 10% glycerol, 1 mM CaCl_2_). A mixture of asolectin lipid (Sigma-Aldrich) and hexadecane was painted across the aperture to establish a planar bilayer membrane as previously described (*47*). To promote liposome fusions, an osmotic gradient across the bilayer was established by perfusion of Buffer C (20 mM HEPES, pH 7.4, 600 mM NaCl, 3.36% glycerol, 1 mM CaCl2) into the *cis* well. A voltage of -180 mV was applied across the membrane by two Ag/AgCl electrodes and the ion current was measured. Liposomes were added to the *cis* well and mixed via pipetting. Sharp transient spikes in currents were interpreted as successful liposome fusions. After observing channel activity, Buffer D was perfused into the *cis* chamber (20 mM HEPES, pH 7.4, 600 mM NaCl, 10% glycerol) to remove free liposomes and a voltage of +180 mV was applied since channel activity was more stable at +180 mV than -180 mV. Ion current was recorded at 50 kHz and downsampled to 50 Hz for analysis. The recorded ion current is divided by the applied voltage to give units of conductance. All-points histograms of conductance were constructed using a bin width of 2 pS and normalized based on probability density (area under the histogram equals 1).

We observed channels in 8 independent experiments and with two independent protein preps. A subset of experiments was run with controls, where we observed ion channels in 2 out of 3 trials after adding both PEEL-1 liposomes and PMPL-1 liposomes, but we did not see channel activity when adding PEEL-1 alone (n=4) nor PMPL-1 alone (n=3). A typical experiment ran as follows: 2 μL of PEEL-1 liposomes and 2 μL of PMPL-1 liposomes were added to the *cis* chamber and mixed via pipetting. Liposome fusions to the bilayer were confirmed by transient spikes in conductance, beginning approximately 5-20 minutes after addition and channel activity was seen approximately 5-60 minutes after fusions began. The amount of time until channels were observed was used as a benchmark for control experiments, where 2 μL of either PEEL-1 liposomes or PMPL-1 liposomes were added to the chamber and allowed to fuse for the same amount of time or longer than experimental runs.

## Acknowledgements

We thank Amy Clippinger, Chau Vuong, Irini Topalidou, Michael Crawford, Tyler Couch, Emma Mackey, Teresa Swanson, Andrew Oberst, Adam Steinbrenner, Sharona Gordon, Bertil Hille, Jihong Bai, Alexey Merz, Suzanne Hoppins, Harmit Malik, and members of the Malik lab (Fred Hutch) for their technical help and input throughout this project. We thank Aaron Severson for the AFS216 strain and Jérôme Cattin-Ortolá for isolating *yak52*. Some strains were provided by the CGC, which is funded by NIH Office of Research Infrastructure Programs (P40 OD010440). This work was supported by the following grants:

National Institutes of Health Institutional Training Grant T32 GM007270 (LC)

National Science Foundation CAREER Award MCB-1552101 (MA)

National Science Foundation Grant MCB-2344838 (MA)

National Institutes of Health Grant R01 HL144801 (J-MR)

National Institutes of Health Grant R01 HL151389 (J-MR)

National Institutes of Health Grant R01 HL126523 (J-MR)

National Human Genome Research Institute R01HG005115 (AHL, JHG)

## Author contributions

Conceptualization: LC, ADW, CAT, GP, MA

Data curation: LC, ADW, CAT, GP, MCF, SJA

Formal Analysis: LC, ADW, CAT, GP

Funding acquisition: LC, AHL, JHG, J-MR, MA

Investigation: LC, ADW, CAT, GP, AU, MCF, SJA, MA

Methodology: LC, ADW, CAT, GP, AU, AHL, JHG, MA

Project administration: LC, MA

Resources: LC, ADW, CAT, GP, AU, MCF, SJA, AHL, JHG, J-MR, MA

Software: LC, ADW, CAT, GP

Supervision: AHL, JHG, J-MR, MA

Validation: LC, ADW, CAT, GP, AU, MCF, SJA

Visualization: LC, ADW, CAT

Writing – original draft: LC, ADW, CAT

Writing – review & editing: LC, ADW, CAT, AHL, MA

## Competing interests

Authors declare no competing interests.

## Data and materials availability

All data are available in the manuscript or the supplementary materials.

## Supplementary Materials

Figs. S1 to S16

Tables S1 to S2

Movie S1

Data S1

## Notes

### Competing Interest Statement

The authors have declared no competing interest.

