## Supplemental figures and tables for "Mechanism of an animal toxin-antidote system"

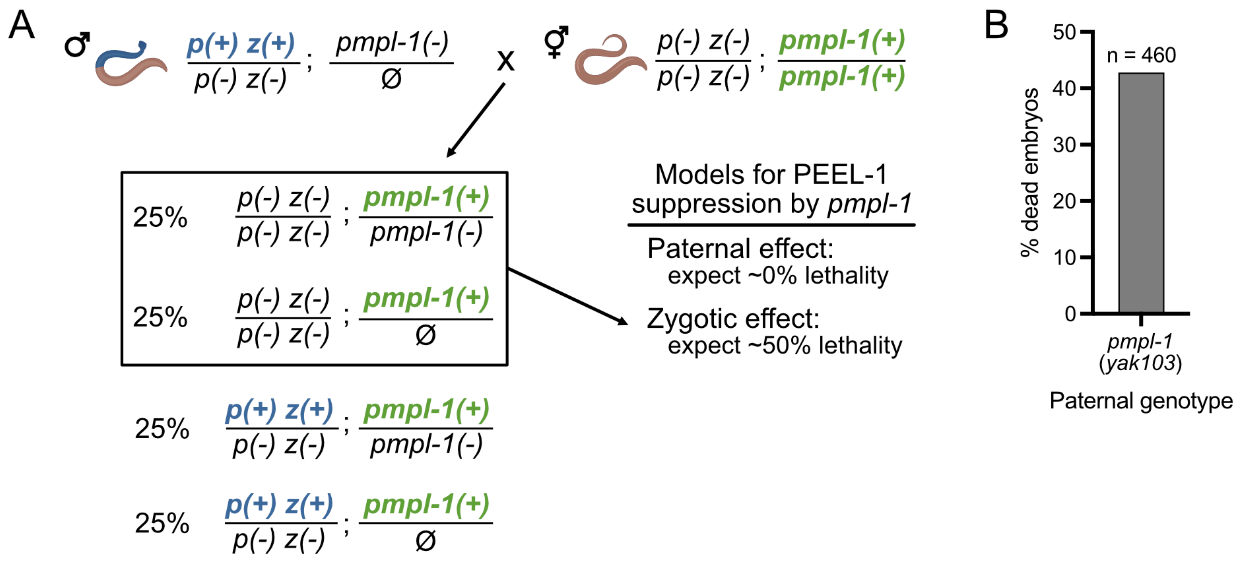


**Fig. S1.** *pmpl-1*(*yak103*) does not act through paternal-effect.

**(A)** Genetic cross to test for paternal versus zygotic effect of sperm-delivered PEEL-1 suppression by *pmpl-1(yak103)* deletion mutant (denoted *pmpl-1(-)*). Males heterozygous for the selfish element and carrying *pmpl-1(-)* are mated to *p(-) z(-); pmpl-1(+)* hermaphrodites. Suppression via paternal effect would result in ~0% embryonic lethality while suppression by a zygotic effect would result in ~50% lethality. **(B)** The percent of dead embryos seen from the cross shown in panel (A). Results are consistent with a model of *pmpl-1*(*yak103*) suppression of PEEL-1 through zygotic effect.

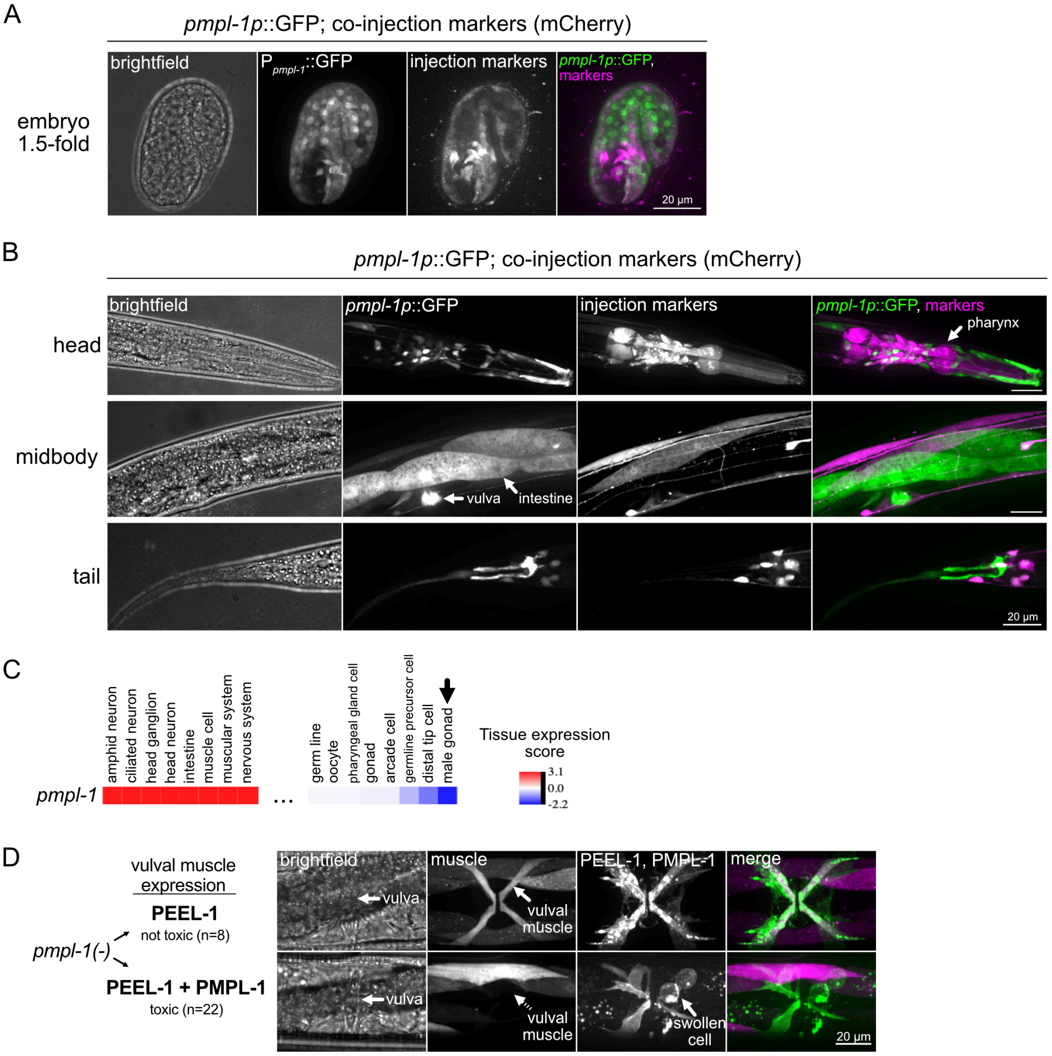


**Fig. S2.** *pmpl-1* expression pattern in *C. elegans*.

**(A)** Maximum intensity projection of a *C. elegans*, 1.5-fold embryo with GFP driven by the *pmpl-1* promoter and co-injection markers *myo-2p*::mCherry, *rab-3p*::mCherry, and *myo-3p*::mCherry. Toxicity from sperm-delivered PEEL-1 occurs after this embryonic stage. **(B)** Maximum intensity projections of hermaphrodite adult head (top), midbody (middle), and tail (bottom) of the same strain shown in panel (A). **(C)** Heatmap of *pmpl-1* of tissue expression scores from RNA-seq dataset of Day 1 adult worms from Kaletsky et al., 2018 (*6*). Tissues with the highest *pmpl-1* expression (red) and lowest expression (blue) are shown. Only the 8 most highly expressed tissues (left) and the 8 most lowly expressed tissues (right) are shown. Expression of *pmpl-1* is lowest in the male gonad (arrow). **(D)** *pmpl-1(yak103)* worms with vulval muscle cell expression of *peel-1*::GFP alone (top) or with *pmpl*-1::GFP (bottom). Number of scored worms are indicated. Green channel brightness is increased in the bottom panel to show fluorescence in the vulval muscle, since toxicity in this cell likely caused decreased levels of fluorescent-tagged proteins. The vulval muscle appears swollen in the green channel when co-expressing *peel-1* and *pmpl-1*. This is the same worm as shown in Figure 1E.

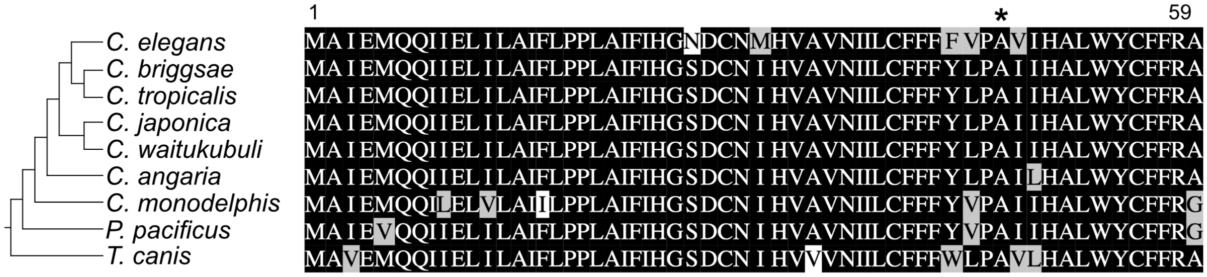


**Fig. S3.** PMPL-1 is conserved in nematodes.

Alignment of PMPL-1 amino acid sequences (right) between representative *Caenorhabditis* species, *Pristionchus pacificus*, and *Toxocara canis* shown in a species phylogeny (left). The *pmpl-1* (*yak52*) allele is mutated at conserved residue A47 (*).

```
PMPL-1 (F47B7.1)     1  ------------------------------MAIEMQQIIELILAIFLPPLAIFIHGNDCNMH
PMPL-2 (C04G6.5)     1  ------------------------------MATDADVIIEVILCIFLPPLAIWWHTKECDIN
PMPL-3 (F25H5.8)     1  -----------------------------MAETPEDKIVMVLLILLFPPLAVWYKEKTCGVG
PMPL-4 (Y55F3BL.6)   1  ----------------------------------MCTILQVIFAFLFPPISVLLT-SGCGLH
PMPL-5 (T23F2.3)     1  ------------------------------MALTCTDIPKFICAVLLPPIGVFLE-KGCDYH
PMPL-6 (T23F2.4)     1  ------------------------------MAITCMDIPKFLFALLLPPVGVFLE-KGCTHH
PMPL-7 (T23F2.5)     1  ------------------------------MALTCTDIPKFLCALLLPPIGVWLE-KGCTYH
PMPL-8 (W02A2.9)     1  ------------------------------MPITCTDIPKFICALLLPPIGVWME-KGCGAD
PMPL-9 (ZK632.10)    1  ----------------------------------MCQILLAILAIFLPPIAVLLD-VGCNCD
PMPL-10 (W10C8.6)    1  ------------------MTVVNVDTKTGYIETDNDRLIMVILLIFLPPLAVFFKSRGCTSQ
PMPL-11 (R10D12.6)   1  -----------MELSEVSVVSVSEEEAKFYIETDNDRLVMALMWLILPPMAVYFKSRGCTKH
PMPL-12 (R10D12.7)   1  -----------MEMAEVNVVAVPEENRQTYLETDNDRLVMAIIWLIMPPMAVFFKCRGCTKH
PMPL-13 (T06C12.9)   1  -----------MEMSDINCGPSEIEVRNPYIETDNDRLVMVLLMLVLPPMAVYFKGRGCTKH
PMPL-14 (T23B3.2)    1  MAEEKMTANVPADAEEGRVFVVESNRRDEMIKL----VLLIILIVIFPPAAVAVHANECNMH
PMPL-15 (W03G9.10)   1  ---------------------MSQNIPETVEKTDTDLLIMALLLIVFPPLGVLLKSNGFTPP

PMPL-1 (F47B7.1)    33  VAVNIILCFFFFVPAVIHALWYCFFRA----------------------------------
PMPL-2 (C04G6.5)    33  VLIDIIFCLLFWLPGILYAVYICFFRK----------------------------------
PMPL-3 (F25H5.8)    34  VCINVVLYILLIFPAYIHAVYVCYIRDRQ--------------------------------
PMPL-4 (Y55F3BL.6)  28  LLLSILLTCLFVIPGIIHALYLVCCHKH---------------------------------
PMPL-5 (T23F2.3)    32  LAICILLTILGYIPGIIYACYVILAY-----------------------------------
PMPL-6 (T23F2.4)    32  LAICILLTILGYIPGIIYACYIILAY-----------------------------------
PMPL-7 (T23F2.5)    32  LAINILLTILGYIPGIIHACYVILAY-----------------------------------
PMPL-8 (W02A2.9)    32  LVINIVLTILGFIPGVIHACFIICWY-----------------------------------
PMPL-9 (ZK632.10)   28  LLINILLTCLGIIPGIIHAWYIILCKEKTVVQNIYVQTNDHGTAPPAYSPYSA--------
PMPL-10 (W10C8.6)   45  VCLNILLYIFLIIPAYCHATWYCFIRGREHEVRAELSRRI---------------------
PMPL-11 (R10D12.6)  52  VCLNVLLYFFLILPSYIHATWYCFVRGRQCEAEDGFVRAR---------------------
PMPL-12 (R10D12.7)  52  VFINFLLYLLLVLPAYKHATWFCFVKGREFEAEDGFVRAR---------------------
PMPL-13 (T06C12.9)  52  VLINIFLYILLVLPAYKHATWFCFVKGRECEAENGFVRVR---------------------
PMPL-14 (T23B3.2)   59  VFISLILVFFFMIPSYIHAIWYCFFRKPTQMTIS---------------------------
PMPL-15 (W03G9.10)  42  VFISFFLYFLFILPSYIFSVWYCFVQQRKDSILPLSSNDFHNNLALNSISASVSHKDIQVY
```

**Fig. S4.** 15 PMP3-like proteins in *C. elegans*.

Alignment of all 15 PMP3-like proteins in *C. elegans* identified by BLAST.

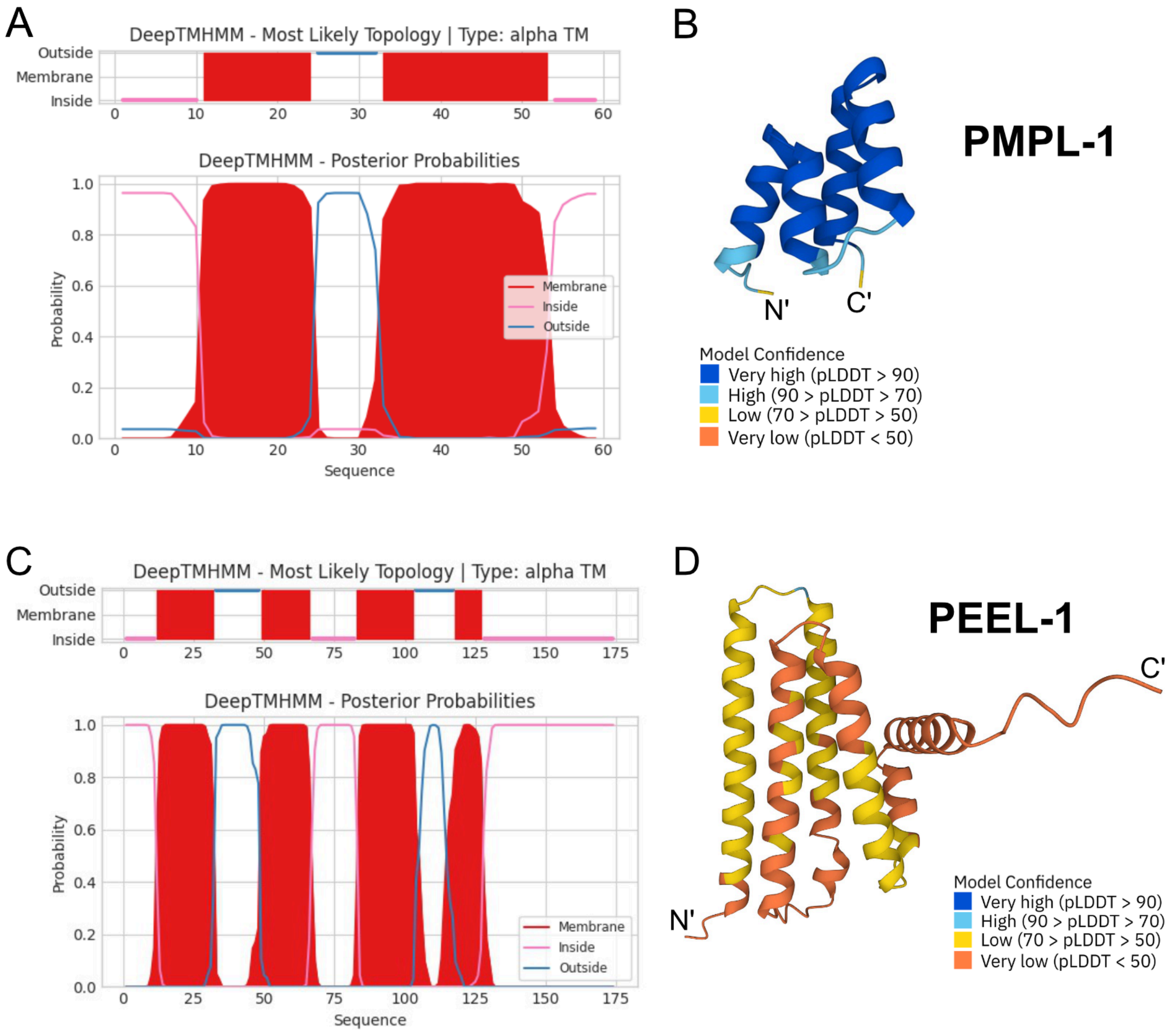


**Fig. S5.** Structural predictions of PEEL-1 and PMPL-1.

DeepTMHMM predictions of **(A)** PMPL-1 and **(C)** PEEL-1. PMPL-1 is predicted to be a two-pass transmembrane protein. PEEL-1 is predicted to be a four-pass transmembrane protein. Both proteins are predicted to have their N- and C-termini facing the cytosol. Structural predictions from AlphaFold2 of **(B)** PMPL-1 and **(D)** PEEL-1. AlphaFold2 predicts a very high-confidence PMPL-1 structure that suggests it is a monotopic protein (passing through one leaflet of a lipid bilayer). The PEEL-1 structure is of low and very-low confidence, likely because there are no known homologs of this protein.

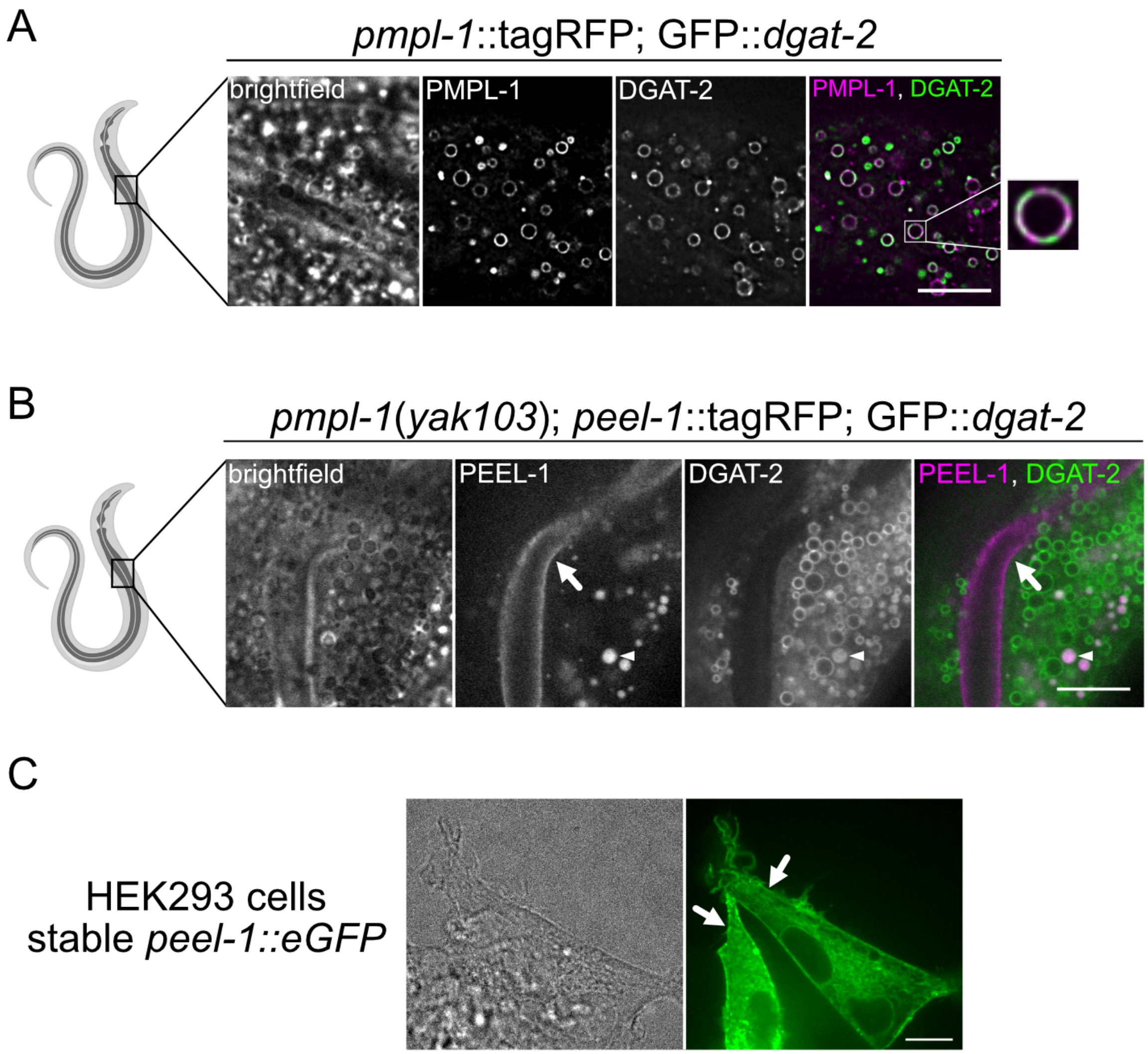


**Fig. S6.** PEEL-1 and PMPL-1 localization.

**(A-B)** Intestinal cells of an adult *C. elegans* worm with the indicated constructs. **(A)** Wild-type worms expressing *pmpl-1*::tagRFP and GFP::*dgat-2*. DGAT-2 localizes to lipid droplet membranes, and PMPL-1::tagRFP co-localizes to these organelles. Inset shows one lipid droplet. **(B)** *pmpl-1*(*yak103*) expressing GFP::*dgat-2* and *peel-1*::tagRFP. PEEL-1 signal (arrow) appears on plasma membrane lining the intestinal lumen and does not co-localize with lipid droplets. Autofluorescence from gut granules appears as filled-in circles in both channels (arrowhead). **(C)** HEK293 cells stably expressing *peel-1*::eGFP. PEEL-1::eGFP localizes to the ER and plasma membrane (arrows). Scale bar = 10µm.

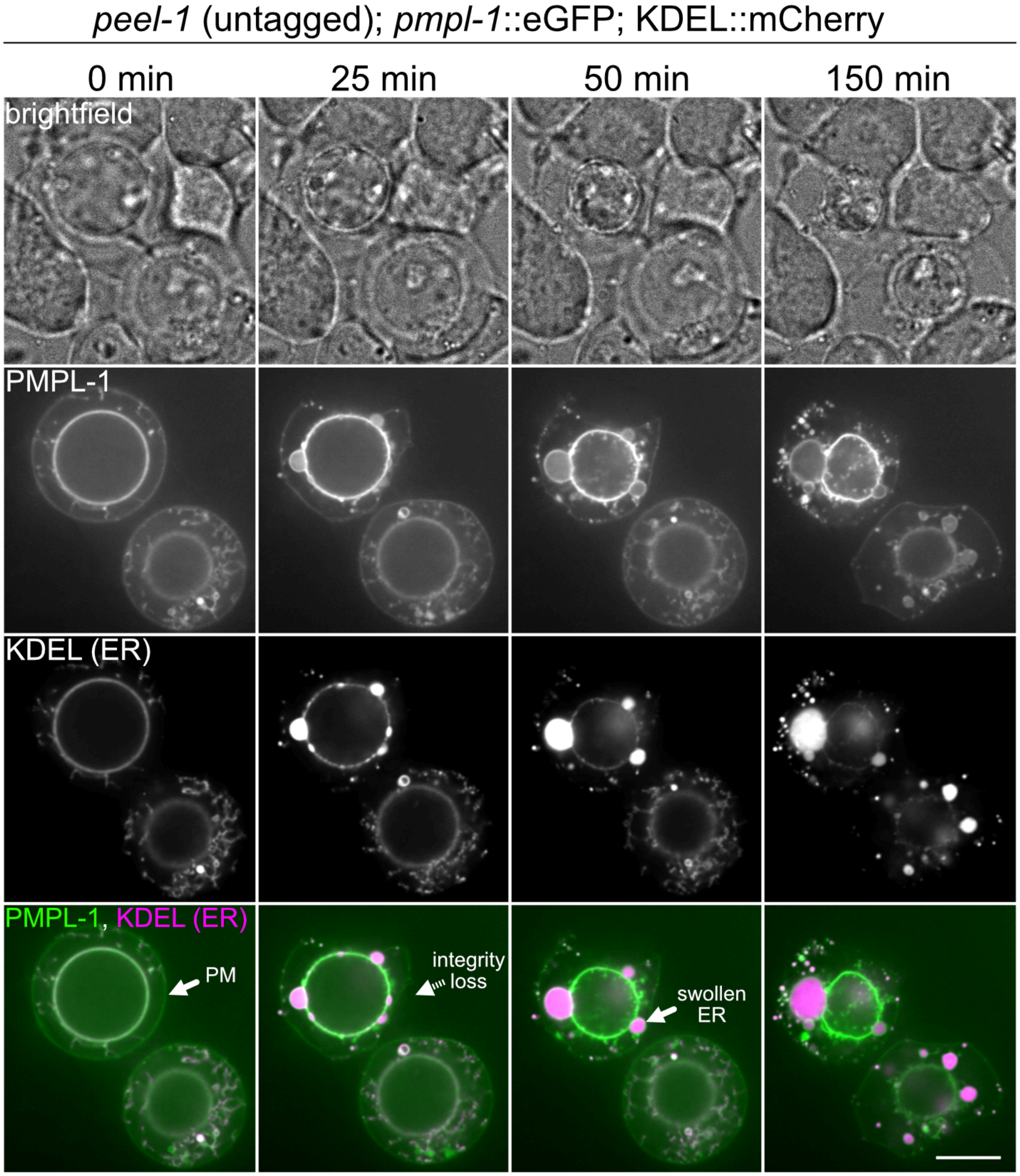


**Fig. S7.** Toxicity causes ER swelling, ER fragmentation, and cell lysis.

Live-cell imaging time course of two HEK293T cells transfected with constructs coding for PEEL-1 (untagged), PMPL-1::eGFP, and mCherry::KDEL (ER marker). Imaging began at 20 hours post-transfection (t=0 min). The top cell experiences a loss of plasma membrane (PM) integrity at 25 min, followed by ER swelling. The lower cell loses PM integrity at 150 min. Scale bar = 10µm.

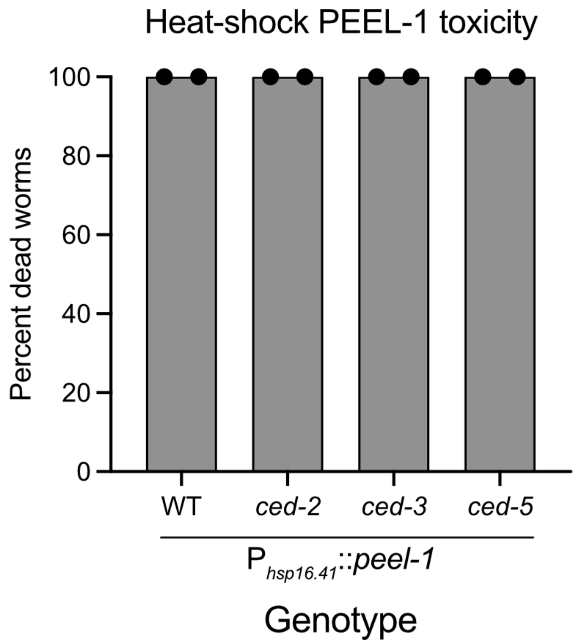


**Fig. S8.** Ectopic PEEL-1 toxicity is non-apoptotic.

Percent dead worms after heat-shock induced expression of PEEL-1. Worms deficient in apoptosis (*ced-3*) and cell engulfment (*ced-2* and *ced-5*) were tested. Two independent experiments were done, with n=50 worms for each data point.

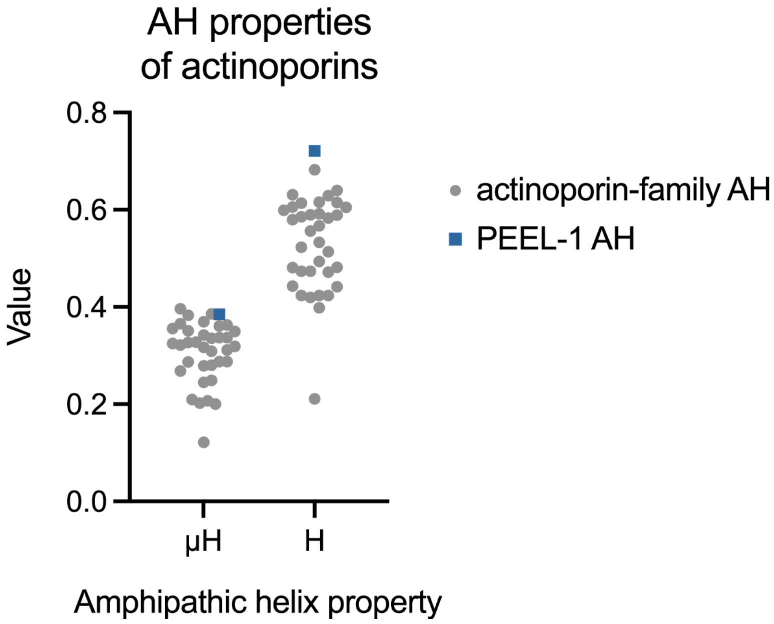


**Fig. S9.** Amphipathic helix properties of Actinoporin toxins.

Hydrophobic moment (μH) and hydrophobicity (H) of the amphipathic helix of 35 Actinoporin toxin proteins (data acquired from Macrander and Daly, 2016 (*29*)) (gray circles) and the predicted PEEL-1 amphipathic helix (blue square).

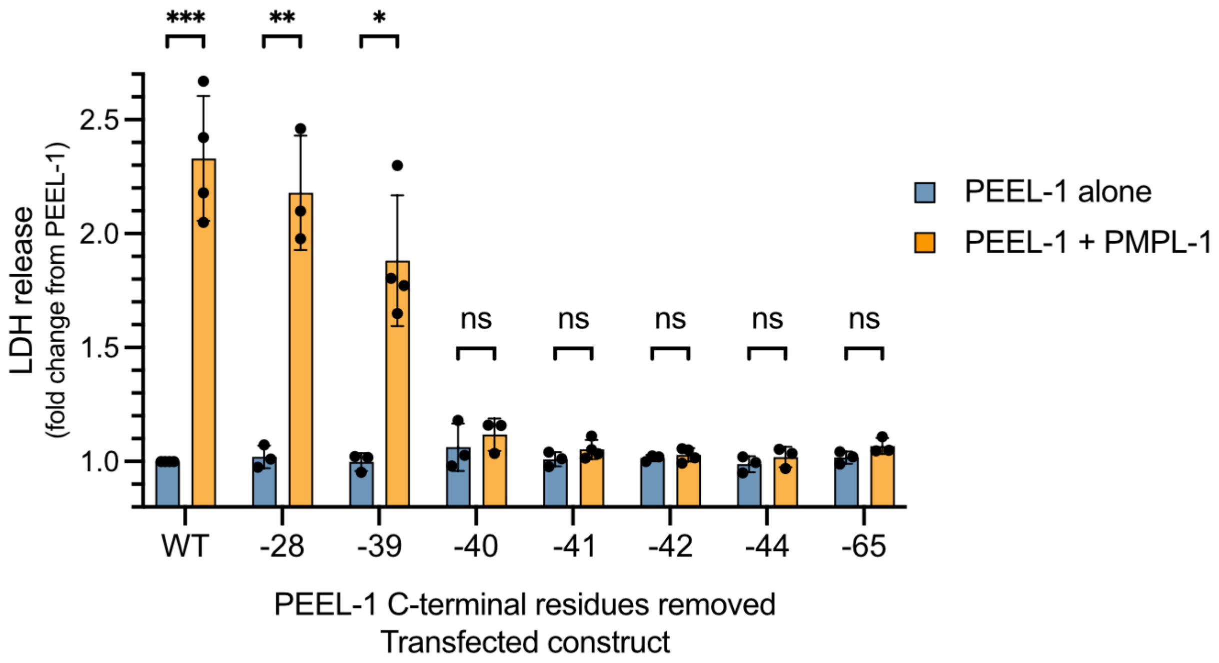


**Fig. S10.** Toxicity of PEEL-1 C-terminal truncations.

Cytotoxicity of PEEL-1 C-terminal truncations assayed alone (blue bars) or with PMPL-1 (yellow bars) in HEK293T cell transfections. The number of amino acids removed from the C-terminus is indicated (ex. -28 means 28 amino acids were removed). Data from PEEL-1 WT, -28, -39, and -65 are from Figure 4C. Toxicity is lost upon removal of the -40 residue (Ala135). Statistical tests done using multiple unpaired t-tests with Holm-Šídák test, comparing each PEEL-1 alone to PEEL-1 and PMPL-1 (*, $p < 0.05$; **, $p < 0.01$; ***, $p < 0.001$).

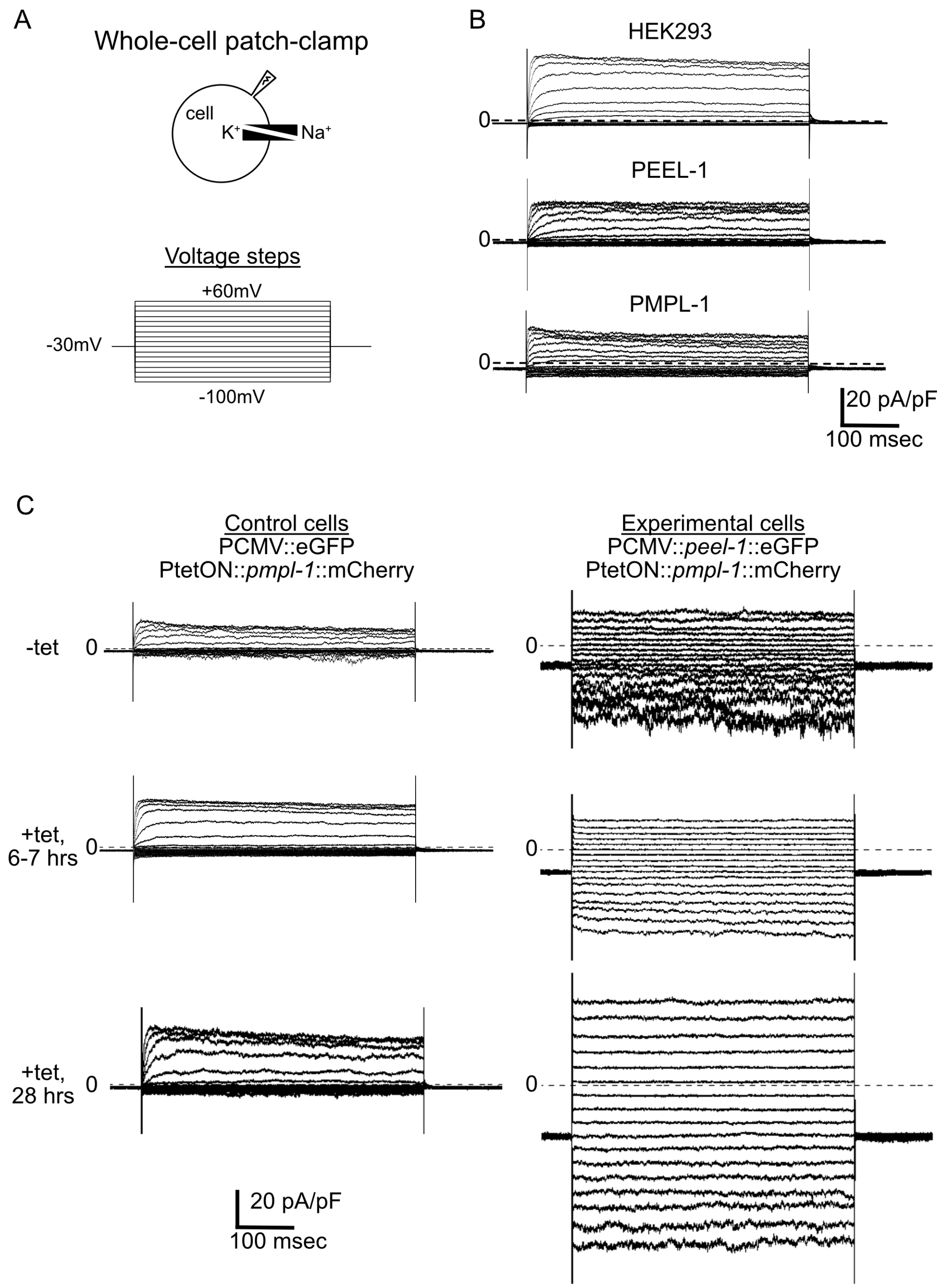


**Fig. S11.** Raw traces from HEK293 electrophysiology experiments.

**(A)** Schematic of electrophysiology experiment. High intracellular potassium (140 mM $K^+$/ 8.6 mM $Na^+$) and high extracellular sodium (145 mM $Na^+$/ 4 mM $K^+$) solutions are used. Currents elicited by a family of 0.5 second voltage steps from a -30 mV holding potential, from -100 mV to 60 mV, in 10 mV increments. Current traces are normalized to capacitance (pF) and not leak-subtracted. **(B)** Representative traces from untransfected HEK293 cells (top) and HEK293 cells transfected with *peel-1*::eGFP (middle) or *pmpl-1*::mCherry (bottom). Scale bar shown (bottom-right). **(C)** Representative traces of tetracycline-inducible cells lines. Control cells (left) have stable expression of eGFP and inducible expression of *pmpl-1*::mCherry. Experimental cells (right) have stable expression of *peel-1*::eGFP and inducible expression of *pmpl-1*::mCherry. Recordings of cell lines without induction (top), 6-7 hours after tetracycline addition (middle), and 28 hours after tetracycline addition (bottom) are shown.

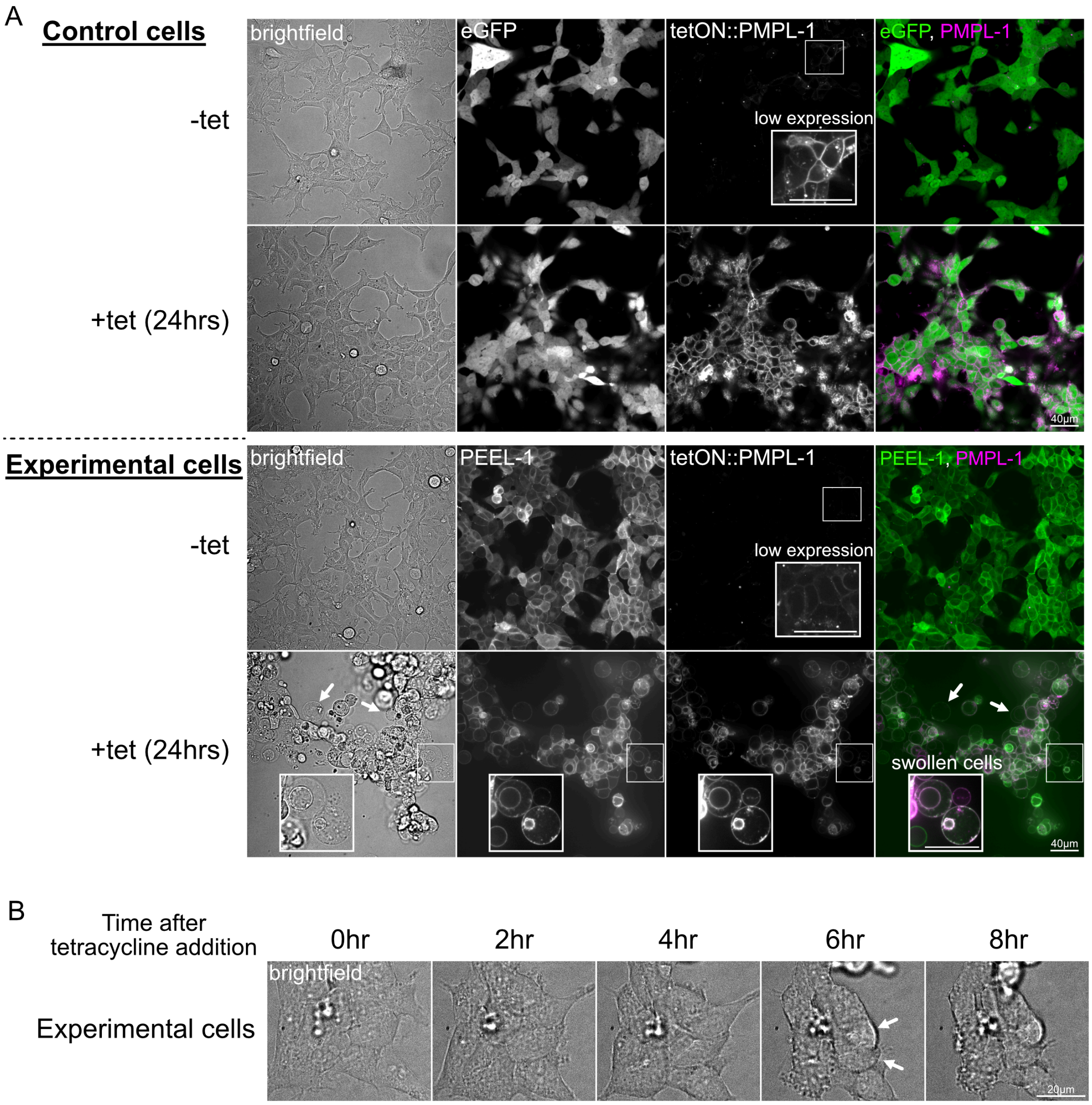


**Fig. S12.** Tetracycline-induced toxicity in stable cell lines.

**(A)** Live-cell images of Control cells (CMV-driven eGFP; tetON::*pmpl-1*::mCherry) and Experimental cells (CMV-driven *peel-1*::eGFP; tetON::*pmpl-1*::mCherry). Cell lines are shown without tetracycline (-tet) and 24 hours after tetracycline addition (+tet (24hrs)). Insets in -tet conditions show leaky expression of PMPL-1 in both cell lines (LUT adjusted within inset). Images show successful tetracycline-inducible expression of PMPL-1::mCherry and efficient tetracycline-induced killing in experimental cells but not control cells. Arrows and inset in experimental cells +tet (24 hrs) show examples of swollen cells. Exposure time in the green channel is different between cell lines. Scale bar = 40 µm. **(B)** Time course of toxicity after addition of tetracycline to experimental cells. Noticeable cell swelling is seen after 6 hours (arrows). Some acute swelling may also be visible at 4 hours after addition of tetracycline. Scale bar = 20 µm.

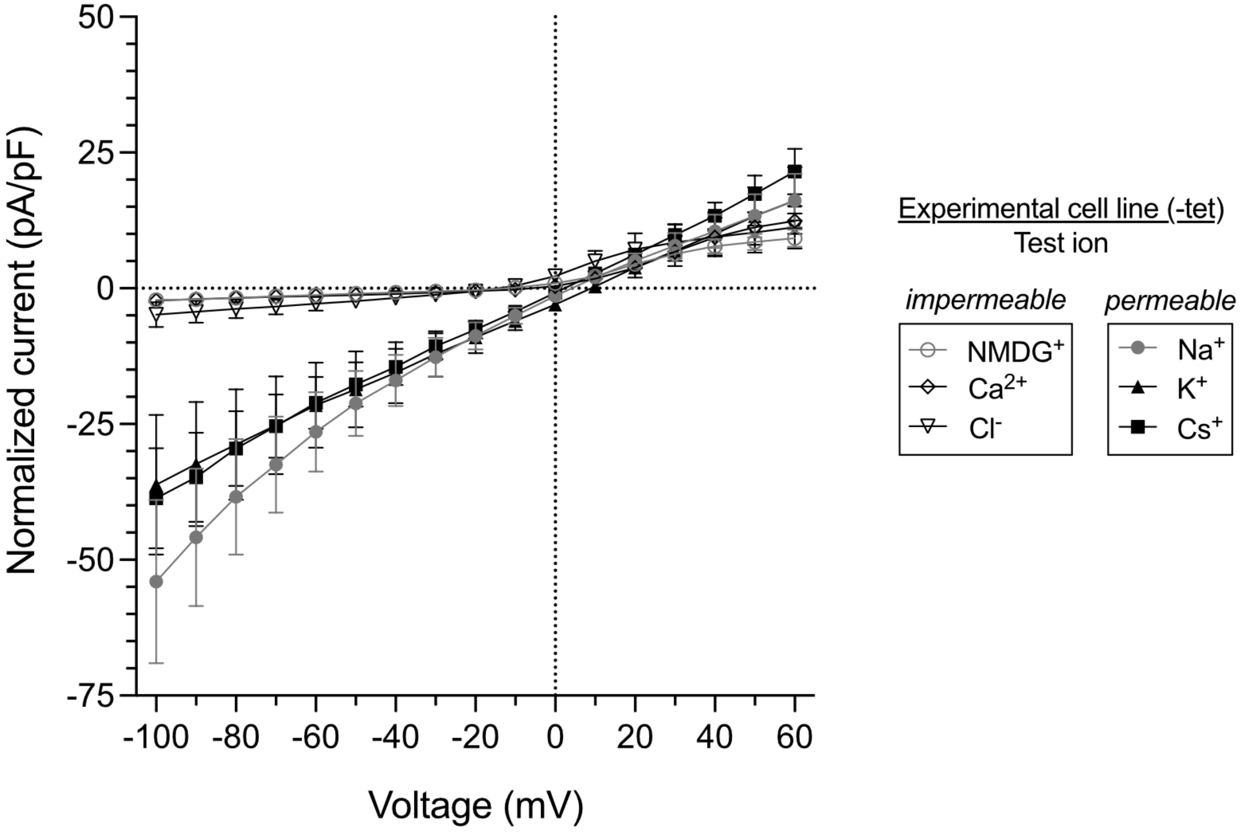


**Fig. S13.** Uninduced Experimental cells are permeable to monovalent cations.

Current-voltage plots of Experimental cells without tetracycline in different ionic conditions to test permeabilities of the indicated ions. Permeable ions have greater inward currents ($Na^+$, $K^+$, $Cs^+$; closed symbols) compared to impermeable ions ($NMDG^+$, $Ca^{2+}$, $Cl^-$; open symbols) at negative voltages. Mean with SEM is shown. These data are summarized in Fig. 5E.

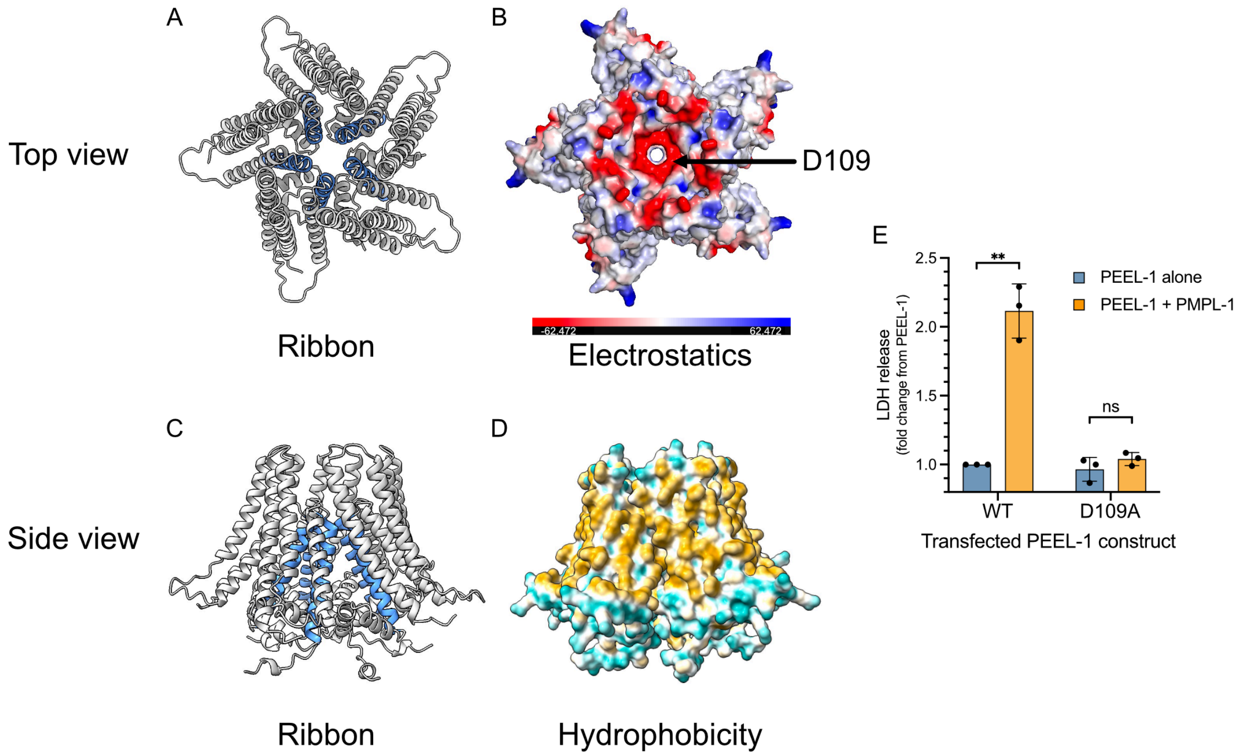


**Fig. S14.** Predicted PEEL-1 pentameric structure.

AlphaFold2 prediction of the PEEL-1 pentameric structure is shown. Two angles are shown: **(A-B)** top view, showing the predicted extracellular face of the complex, and **(C-D)** side view, in the plane of the lipid bilayer. **(A and C)** Ribbon diagram with the amphipathic helix colored in blue creating the lining of a pore-like region. **(B)** Surface representation of electrostatic predictions (red = negative charge, blue = positive charge). An uninterrupted hole can be seen through the structure, with a ring of negative charge from five D109 residues. **(D)** Surface representation of hydrophobicity (yellow = hydrophobic, cyan = hydrophilic). **(E)** Cytotoxicity of mutants which eliminate the predicted ring of negative charge at the top of the complex via a D109A mutation. Plot shows mean with SD. Statistics performed using multiple unpaired t-tests with Holm-Šídák test. All tested comparisons are shown. PDB file available in data S1.

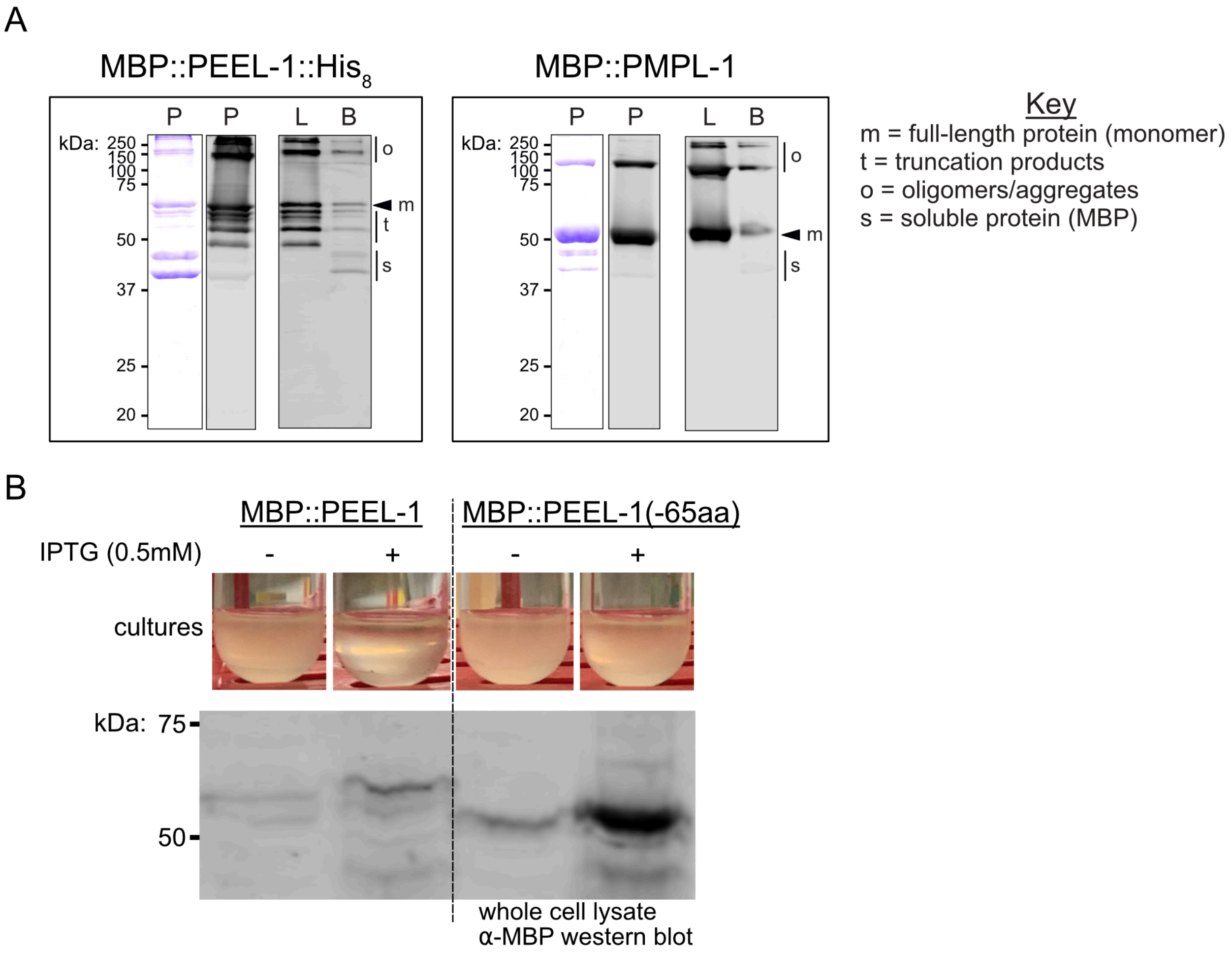


**Fig. S15.** PEEL-1 and PMPL-1 purification and liposomes.

**(A)** SDS-PAGE gels of indicated constructs after purification (P; Coomassie Blue stain and anti-MBP western blot) and after incorporation into liposomes (L; anti-MBP western blots). Liposomes were collected after flotation through a density gradient before use in synthetic bilayer experiments. The top fraction contains liposomes (L) and the bottom fraction (B) contains unincorporated protein. Monomers (m), oligomers or aggregates (o), and truncation products of PEEL-1 (t) are indicated. The bottom bands seen in both MBP-tagged PEEL-1 and PMPL-1 purifications are soluble proteins (s) that did not get incorporated into liposomes, likely endogenous MBP (43 kDa) or truncated MBP-tagged protein. **(B)** PEEL-1 is toxic to bacteria and toxicity requires the amphipathic helix. *E. coli* C41(DE3) cells with constructs encoding IPTG-inducible expression of either MBP::PEEL-1 (left) or MBP::PEEL-1(-65aa) (right). Images of cultures are taken after shaking at 18°C overnight, with or without 0.5 mM IPTG. Cultures appear lysed when full-length PEEL-1 is expressed but not PEEL-1(-65aa) which lacks the PEEL-1 AH. Western blot (anti-MBP) of corresponding whole-cell lysates are shown (bottom), confirming higher expression of MBP::PEEL-1(-65aa). This experiment was repeated four times and yielded similar results.

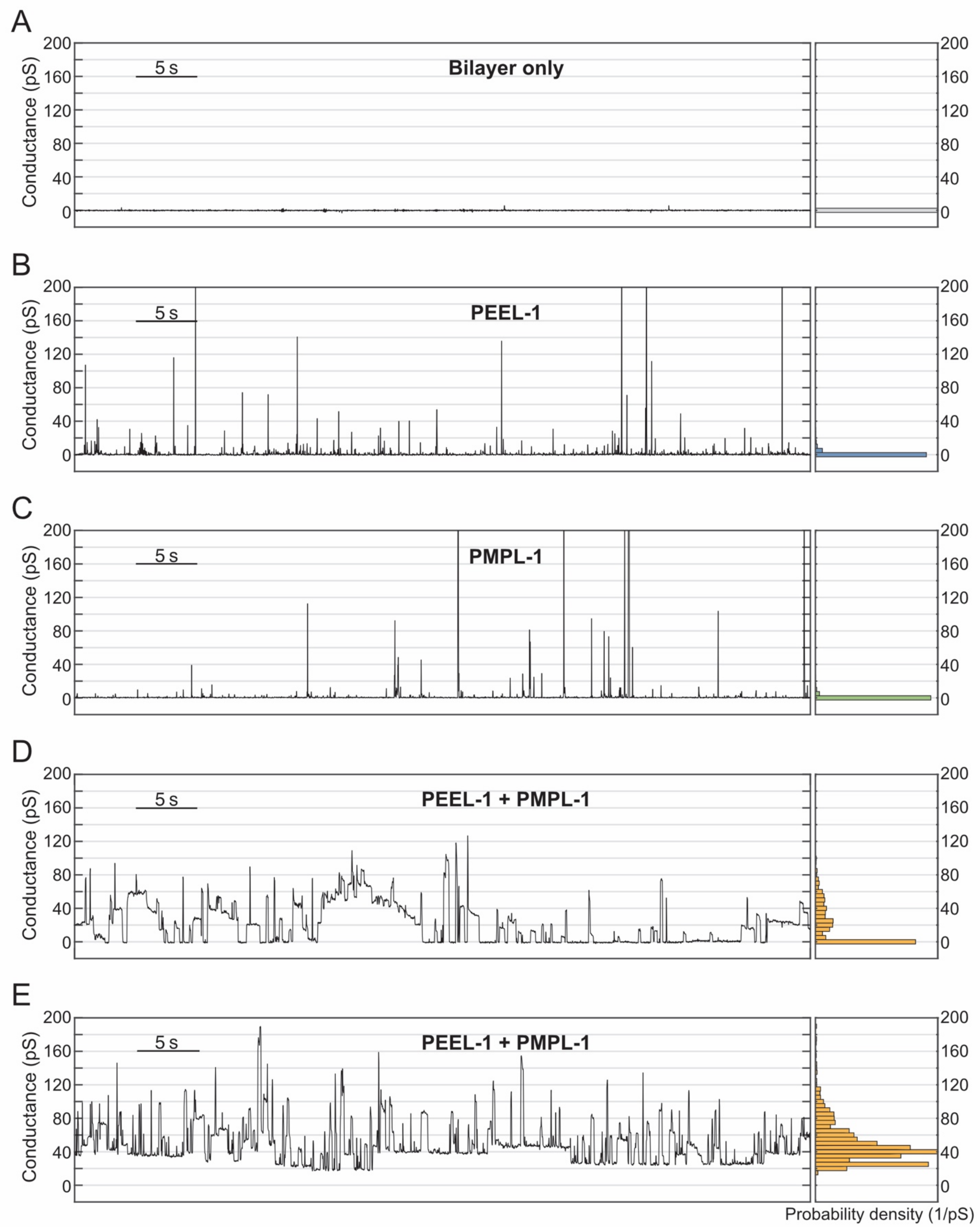


**Fig. S16.** Purified PEEL-1 and PMPL-1 conduct ions through planar lipid bilayers.

Conductance traces through artificial planar lipid bilayers are shown. **(A)** Bilayer alone without addition of liposomes, **(B)** bilayer with PEEL-1 liposomes added (transient spikes indicate successful liposome fusions), **(C)** bilayer with PMPL-1 liposomes added, and **(D-E)** two independent experiments of bilayers with PEEL-1 and PMPL-1 liposomes added. An all-point histogram is shown for each trace (right, 2 pS bin width, normalized based on probability density). Channel activity after addition of PEEL-1 and PMPL-1 was observed in 8 independent experiments. A voltage of -180 mV was applied to the bilayer for liposome fusions in all experiments. After observing channel activity, a voltage of +180 mV was applied since channel activity was more stable at positive voltages. Bottom two panels show traces at +180 mV. Scale bar = 5 seconds.

**Table S1.** *C. elegans* strains used in this study.

| Strain ID | Strain genotype | Source |
|---|---|---|
| N2 | N2 | CGC |
| XZ1372 | yakTi4[hsp-16.41p::eGFP::his-44 , NeoR] I ; oxSi507[hsp-16.41p::peel-1, Cb-unc-119] II ; oxSi280[hsp-16.41p::peel-1, Cb-unc-119] IV | Crawford et al., 2023 |
| XZ1047 | oxSi507[hsp-16.41p::peel-1, Cb-unc-119] II ; unc-119(ed9) III ; oxSi280[hsp-16.41p::peel-1, Cb-unc-119] IV ; him-5(e1490) V | Crawford et al., 2023 |
| XZ103 | oxSi507[hsp-16.41p::peel-1, Cb-unc-119] II ; oxSi280[hsp-16.41p::peel-1, Cb-unc-119] IV ; pmpl-1(yak103) X | this study |
| XZ2283 | oxSi507[hsp-16.41p::peel-1, Cb-unc-119] II ; oxSi280[hsp-16.41p::peel-1, Cb-unc-119] IV ; F47B7.1(yak52) X | this study |
| AFS216 | zeel-1(tm3419) I peel-1(cle6) I | Aaron Severson |
| XZ1177 | oxSi507[hsp-16.41p::peel-1, Cb-unc-119] II ; unc-119(ed9) III ; oxSi280[hsp-16.41p::peel-1, Cb-unc-119] IV ; him-5(e1490) V ; pmpl-1(yak52) X | this study |
| XZ1307 | oxSi507[hsp-16.41p::peel-1, Cb-unc-119] II ; oxSi280[hsp-16.41p::peel-1, Cb-unc-119] IV ; him-5(e1490) V ; pmpl-1(yak103) X | this study |
| EG1000 | dpy-5(e61) I ; rol-6(e187) II ; lon-1(e1820) III | Erik M. Jorgensen |
| EG1020 | bli-6(sc16) IV ; dpy-11(e224) V ; lon-2(e678) X | Erik M. Jorgensen |
| EG8040 | oxTi302[Peft-3::mCherry cb-unc-119(+)] I ; oxTi75[Peft-3::GFP::H2B::tbb-2utr unc-18(+)] II ; oxTi411[Peft-3::TdTomato::H2B::unc-54 cb-unc-119(+)] III ; him-8(e1489) IV | Jorgensen lab |
| EG8041 | oxTi76[Peft-3::GFP::H2B::tbb-2utr unc-18(+)] IV ; oxTi405[Peft-3::TdTomato::H2B::unc-54 cb-unc-119(+)] V him-5(e1490) V ; oxTi421[Peft-3::mCherry cb-unc-119(+)] X | Jorgensen lab |
| XZ2194 | pmpl-1 (yak103) X | this study |
| XZ2103 | oxSi507[hsp-16.41p::peel-1, Cb-unc-119] II ; ced-3(n717) IV ; him-5(e1490) V | this study |
| XZ2102 | oxSi507[hsp-16.41p::peel-1, Cb-unc-119] II ; ced-5(n1812) IV | this study |
| XZ2096 | oxSi507[hsp-16.41p::peel-1, Cb-unc-119] II ced-2(n1994) IV | this study |
| XZ2254 | yakEx195[pmpl-1p::GFP; myo-2p::mcherry; myo-3p::mcherry; rab-3p::mCherry] | this study |
| XZ2276 | pmpl-1(yak103) X ; yakEx203[exp-3p::peel-1::GFP, myo-3p::mCherry] | this study |
| XZ2633 | pmpl-1(yak103) X ; yakEx275[exp-3p::peel-1::GFP, exp-3p::pmpl-1::GFP, myo3p::mCherry] | this study |
| XZ2551 | yakEx264[hsp-16.41p::peel-1(-28aa), cc::GFP] | this study |
| XZ2634 | yakEx276[hsp-16.41p::peel-1(-39aa), cc::GFP] | this study |
| XZ2548 | yakEx263[hsp-16.41p::peel-1(-65aa), cc::GFP] | this study |
| XZ2454 | pmpl-1(yak103) X ; hjSi56[Pvha-6::3xFLAG::TEV::GFP::dgat-2::let-858 3' UTR] IV ; yakEx243[vha-6p::peel-1::tagRFP, myo-2p::mCherry] | this study |
| XZ2452 | hjSi56[Pvha-6::3xFLAG::TEV::GFP::dgat-2::let-858 3' UTR] IV ; yakEx242[vha-6p::pmpl-1::tagRFP, myo-2p::mCherry] | this study |

**Table S2.** Constructs used in this study.

| Construct ID | Description | resistance | sequenced? | notes | | for expression i[illegible] |
|---|---|---|---|---|---|---|
| pLC4 | exp-3p::pmpl-1::tagRFP::tbb-2 3'UTR | carb | | | | worm |
| pLC6 | exp-3p::peel-1::GFP::tbb-2 3'UTR | carb | | | | worm |
| pLC26 | MBP::TEV::peel-1 | carb | sequenced | | | bacteria |
| pLC28 | MBP::TEV::pmpl-1 | carb | sequenced | | | bacteria |
| pLC31 | exp-3p::pmpl-1::GFP tbb-2 3'UTR | carb | | | | worm |
| pLC37 | PMP3(S. cerevisiae)::mCherry_N1 | kan | sequenced | | | mammalian |
| pLC38 | mCherry::zeel-1_N1 | kan | | | | mammalian |
| pLC54 | pmpl-1::eGFP_N1 | kan | | | | mammalian |
| pLC65 | MBP::TEV::peel-1 | carb+chlor | sequenced | | | bacteria |
| pLC67 | MBP::TEV::pmpl-1 | carb+chlor | sequenced | | | bacteria |
| pLC79 | tetON::pmpl-1::mCherry_pFTSH | carb | sequenced | | | mammalian |
| pLC84 | exp-3p::pmpl-1::tagRFP::tbb-2 3'UTR | carb | | | | worm |
| pLC103 | vha-6p::pmpl-1::tagRFP::tb-2 3'UTR | carb | | | | worm |
| pLC113 | vha-6p::peel-1::tagRFP::tbb-2 3'UTR | carb | | | | worm |
| pLC122 | pmpl-2::mCherry_N1 | kan | sequenced | | | mammalian |
| pLC123 | pmpl-1(A47T)::mCherry_N1 | kan | sequenced | | | mammalian |
| pLC124 | peel-1(S124F)::eGFP_N1 | kan | sequenced | | | mammalian |
| pLC172 | hsp-16.41p::peel-1(-28aa)::let-858 3'UTR | carb | sequenced | | | worm |
| pLC173 | hsp-16.41p::peel-1(-65aa)::let-858 3'UTR | carb | sequenced | | | worm |
| pLC174 | peel-1(-28aa)_N1 | kan | sequenced | | | mammalian |
| pLC175 | peel-1(-65aa)_N1 | kan | sequenced | | | mammalian |
| pLC226 | peel-1(-39aa)_N1 | kan | sequenced | | | mammalian |
| pLC227 | MBP::TEV::peel-1(-65aa) | carb | sequenced | | | bacteria |
| pLC294 | MBP::TEV::peel-1 (-65aa) | carb/chlor | sequenced | | | bacteria |
| pLC297 | peel-1(-42aa)_N1 | kan | sequenced | | | mammalian |
| pLC298 | peel-1(-44aa)_N1 | kan | sequenced | | | mammalian |
| pLC304 | peel-1(-41aa)_N1 | kan | sequenced | | | mammalian |
| pLC305 | peel-1(-40aa)_N1 | kan | sequenced | | | mammalian |
| pLC345 | peel-1::eGFP::ER-ret(GBR1 C-tail)_N1 | kan | sequenced | | | mammalian |
| pLC363 | peel-1(S124V)::eGFP_N1 | kan | sequenced | | | mammalian |
| pLC365 | pmpl-1::mCherry::ER-ret(GBR1 C-tail)_N1 | kan | sequenced | | | mammalian |
| pLC370 | MBP::TEV::peel-1::8X His | carb | sequenced | | | bacteria |
| pLC376 | MBP::TEV::peel-1::8X His | carb/chlor | sequenced | | | bacteria |
| pLC385 | peel-1(D109A)::eGFP_N1 | kan | sequenced | | | mammalian |
| pLC395 | peel-1(L115Q)::eGFP_N1 | kan | sequenced | | | mammalian |
| pLC396 | peel-1(L118Q)::eGFP_N1 | kan | sequenced | | | mammalian |
| pLC397 | peel-1(L122Q)::eGFP_N1 | kan | sequenced | | | mammalian |
| pLC398 | peel-1(L126Q)::eGFP_N1 | kan | sequenced | | | mammalian |
| pLC438 | peel-1(L118Q,S124V)::eGFP_N1 | kan | sequenced | | | mammalian |
| pLC439 | peel-1(L118Q,L126Q)::eGFP_N1 | kan | sequenced | | | mammalian |
| pLC440 | peel-1(S124V,L126Q)::eGFP_N1 | kan | sequenced | | | mammalian |
| pLC475 | hsp-16.41p::peel-1(-39aa)::let-858 3'UTR | carb | sequenced | | | mammalian |
| mCherry-KDEL | mCherry-KDEL | kan | | gift from Suzanne Hoppins | | mammalian |
| pGP9 | peel-1::eGFP_N1 | kan | sequenced | | | mammalian |
| pGP10 | pmpl-1::mCherry_N1 | kan | sequenced | | | mammalian |
| pPD97/98 | cc::GFP (unc-122p::GFP) | carb | | gift from Piali Sengupta | | worm |
| pCFJ90 | Pmyo-2::mCherry::unc-54 3'UTR | carb | | | | worm |
| pCFJ104 | Pmyo-3::mCherry::unc-54 3'UTR | carb | | | | worm |
| pGH8 | Prab-3::mCherry::unc-54 3'UTR | carb | | | | worm |
| pBS_SK | pBluescript | carb | | | | worm |
| pCFJ150 | destination vector (4-1-2-3) | carb | | | | |
| eGFP_N1 | eGFP_N1 | kan | | gift from Suzanne Hoppins | | mammalian |
| mCherry_N1 | mCherry_N1 | kan | | gift from Suzanne Hoppins | | mammalian |
| pOG44 | FLP recombinase | carb | | gift from Nancy Maizels | | mammalian |
| pFTSH | vector backbone, tetON expression | carb | | gift from Nancy Maizels | | mammalian |

**Movie S1.** A single HEK293T cell co-expressing PEEL-1::eGFP and PMPL-1::mCherry. Movie starts at 18.5 hours after transfection. One frame = 5 min. Scale bar = 10 µm. Selected frames of this movie are shown in Fig. 3C.

**Data S1.** PDB file of AlphaFold2 prediction of the PEEL-1 pentamer.
